# SoftHybrid: A Hybrid Imputation Algorithm Optimised for Single-Cell Proteomics Data

**DOI:** 10.64898/2026.01.13.699212

**Authors:** Yixin Shi, Simon Davis, Philip D. Charles, Stephen Taylor, Eszter Dombi, Georgina Berridge, Daniel Ebner, Roman Fischer

## Abstract

Missing values (MVs) remain a significant barrier to reliable proteomics analysis, particularly in single-cell proteomics, where small amounts of starting material and limits in detection drive Missing-Not-At-Random (MNAR) sparsity. Existing imputation methods typically target either Missing-At-Random (MAR) or MNAR mechanisms, resulting in a trade-off between replicate consistency and preservation of biological variation, and are largely designed for bulk data. Here, we introduce SoftHybrid, a data-driven imputation framework that jointly models missingness and protein abundance to estimate the probability of MNAR, enabling continuous weighting between MAR- and MNAR-oriented strategies. SoftHybrid requires no external priors (cell type labels, group annotations, predefined missingness assumptions, etc.), enabling fully unsupervised applications. Across ground truth benchmarks and real single-cell proteomics datasets, SoftHybrid outperforms existing methods at low input and matches or exceeds their performance at the mini-bulk level. By preserving proteomic structure and abundance accuracy, it enhances the recovery of biologically meaningful signals. SoftHybrid is implemented as an R package and is freely available on GitHub.

## Introduction

Over the past decade, advances in the sensitivity and throughput of mass spectrometry (MS)- based label-free proteomics have greatly expanded the reach of proteomics studies^1–5^. Both bulk and single-cell approaches now achieve deep proteome coverage, supporting a wide range of applications in basic biology and clinical research^6,7^. Despite this progress, missing values remain a persistent obstacle in proteomics, potentially masking biological effects. Most downstream bioinformatic analyses rely on complete data matrices, and missingness can reduce reproducibility, weaken statistical power, and, most importantly, bias biological interpretation^8–11^. In MS-based proteomics, missingness is a result of complex interactions of technical limitations such as stochastic/incomplete peptide sampling and physicochemical dependencies in complex peptide mixtures (peptide co-elution, ion suppression). Additional effects, such as dynamic exclusion and precursor competition, add further complexity, especially in data-dependent (DDA) acquisition modes^12^. Missing values are generally grouped into two categories: missing not at random (MNAR), arising from limits of quantification, and missing at random (MAR), reflecting stochastic sampling losses^10^. Specifically, MNAR can be further divided into two categories; firstly, proteins whose true biological abundance is close to or below detection limits; secondly, proteins which are biologically absent and normally strongly related to cell types or intervention groups. In practice, however, the ground truth of missingness often remain unknown and real-world datasets usually contain a mixture of both types in varying proportions, shaped by multiple factors such as acquisition strategy, sample complexity, and protein abundance distribution. For instance, the dynamic exclusion and precursor competition inherent in DDA tend to yield more MAR values compared to data-independent acquisition (DIA). This heterogeneity makes missing-value imputation particularly challenging, and remains a requirement for unlocking the potential of quantitative proteomics in biological and medical research.

In practice, most proteomics workflows still rely on a small repertoire of familiar imputation approaches. Methods such as the Gaussian left-censored used in Perseus (Gaussian)^13^, minProb^14^, k-nearest neighbours (KNN)^15^, and random forest (RF)^16^ have become routine choices across the field (Table 1). However, these tools are frequently selected without a clear rationale, often driven more by habit and convention than by addressing the underlying missing-value mechanism. In single-cell proteomics, sparsity is a common feature of acquired data, providing an opportunity to employ missing value imputation methods specifically suitable for this data type. Different studies have tried various solutions, including zero replacement, KNN, minProb, and Gaussian, with Gaussian and KNN being the most frequently used^6,17–20^. However, systematic benchmarking of these methods on single-cell proteomics datasets remains lacking. Vanderaa and Gatto highlighted that, in highly sparse single-cell proteomics data, retaining raw values without imputation may in some cases be preferable^21^, but as, many downstream analyses require a fully observed data matrix, imputation remains a crucial tool to exploit the information obtained by SCP experiments. In general, left-shifted approaches are better suited for missing not at random (MNAR) values, whereas similarity- and structure-based approaches are more effective for missing at random (MAR) values^10^. Yet no single method can address both mechanisms equally well, underscoring the need for hybrid strategies that can be applied consistently across both bulk and single-cell datasets and data acquisition modes^10^.

**Table 1.**
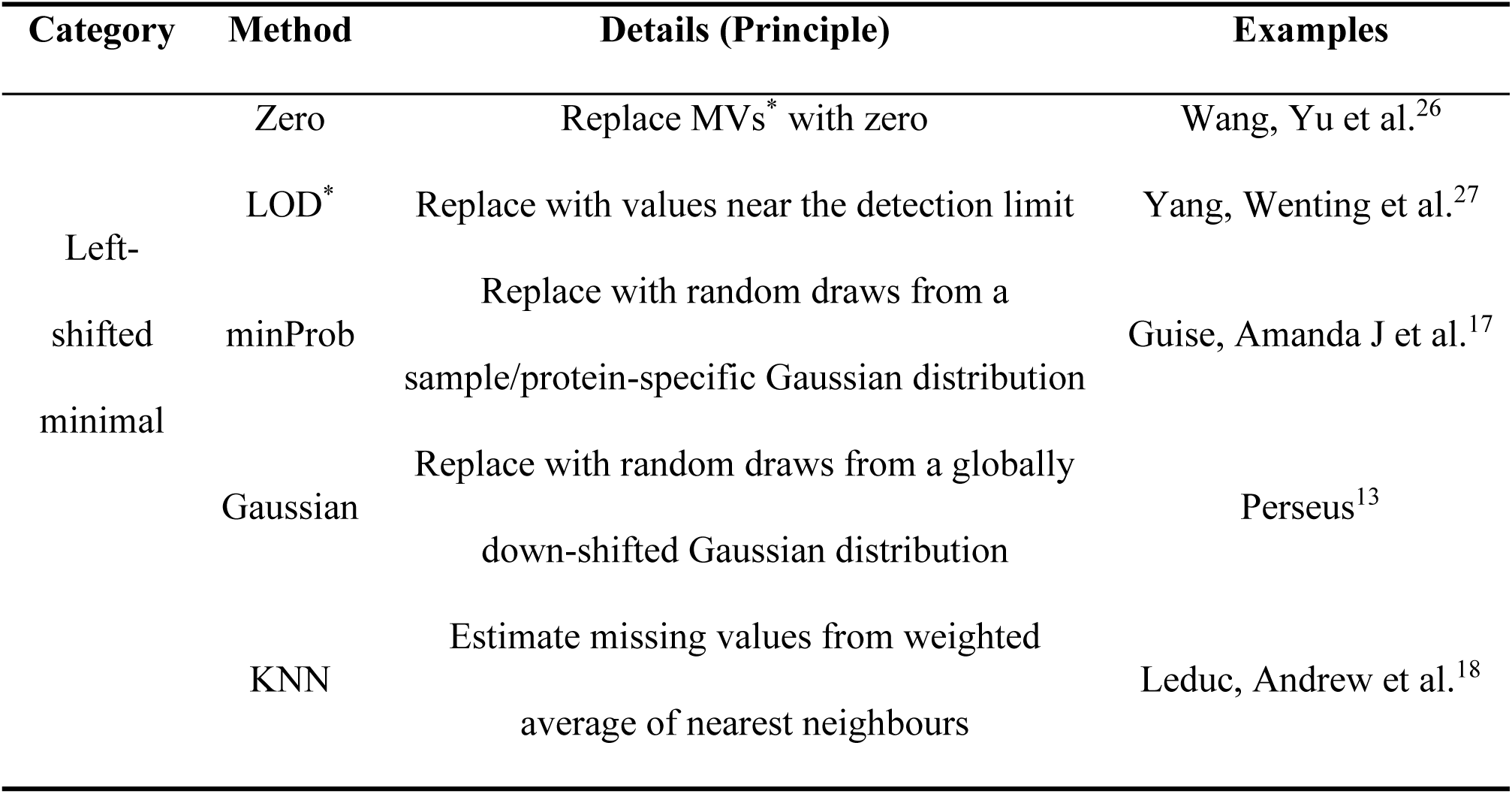
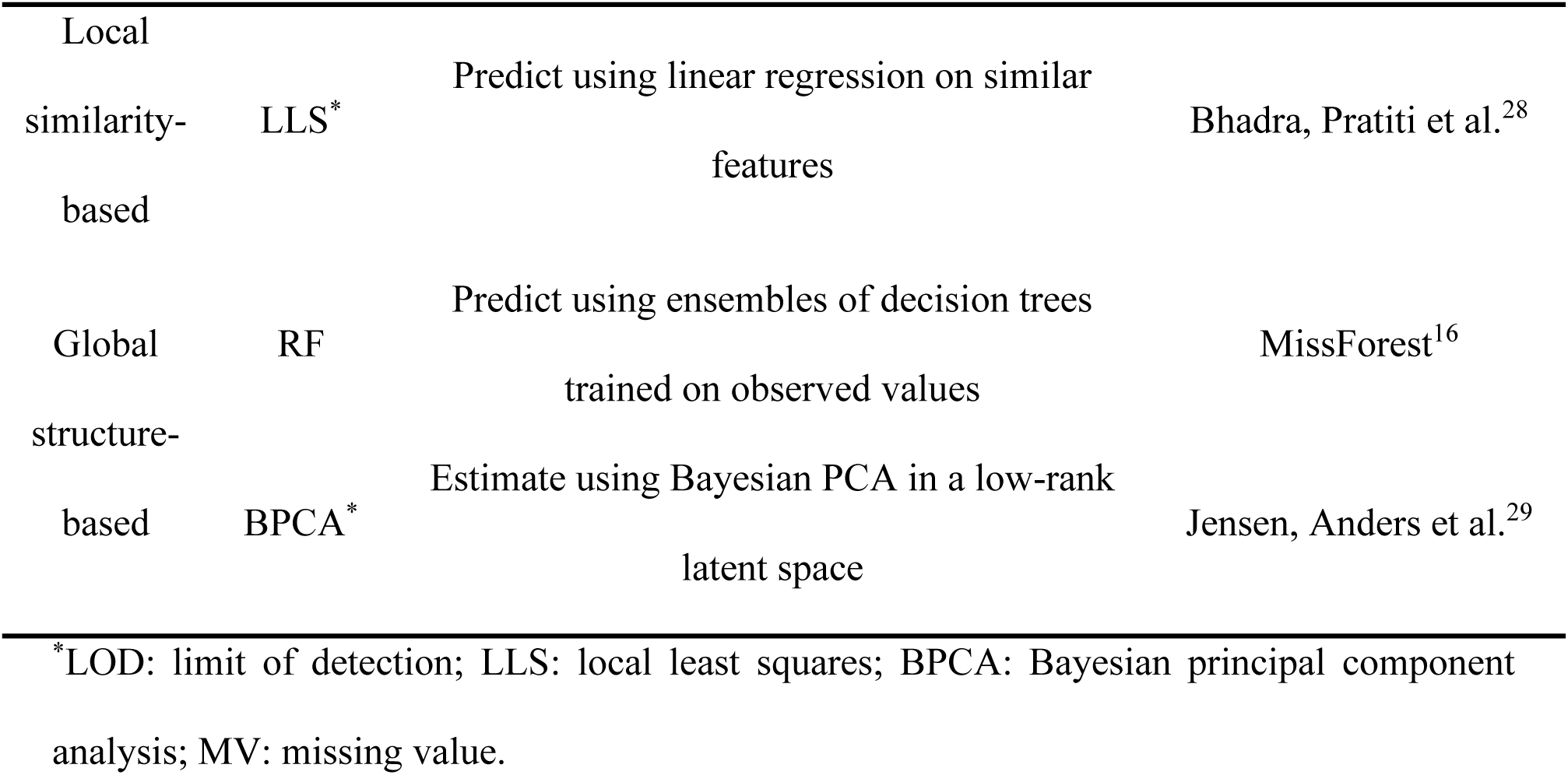
Overview of representative imputation methods and their underlying principles.

Hybrid imputation strategies have emerged in recent years to overcome the limitations of single-method approaches in proteomics. Guo et al. emphasised the biological meaning embedded in missing values and proposed a classifier to distinguish between different missingness mechanisms^22^. However, their work focused on characterising missingness rather than providing an imputation framework. Shi et al. developed a zone-based strategy in which features were partitioned by missing rate and protein abundance, with the best-performing method applied in each zone^23^. While effective, this multi-step design is highly dataset-dependent and difficult to generalise. More recently, MsImpute applied an entropic barycentric distribution to interpolate between MAR- and MNAR-oriented models, achieving better imputation results in bulk datasets with known sample groupings^24^. Its performance in highly sparse single-cell datasets, however, remains uncertain, as the combination of minimal input and extreme sparsity may amplify bias and introduce additional noise into the entropic distribution. Another related modelling strategy has been proposed in the QuantQC framework^25^, where missingness is explicitly modelled by integrating abundance-dependent technical effects with biological information inferred from the data. In this approach, an initial imputation step (e.g. KNN) enables clustering and cell-type annotation, and the resulting labels are subsequently incorporated into regression models to distinguish technical from biological missingness. While this design allows for a more explicit interpretation of missingness mechanisms, it relies on the availability of external information and accuracy of inferred annotations.

Here, we introduce SoftHybrid, an imputation strategy that combines missing rate and protein abundance into a continuous weighting scheme. By smoothly balancing contributions from MAR-oriented and MNAR-oriented models, SoftHybrid avoids the abrupt transitions of zone-based approaches and provides a more flexible framework that adapts to the heterogeneity of real datasets. This design allows it to handle both bulk and single-cell proteomics data as well as DDA and DIA acquisition approaches, while conventional methods often struggle with either over-smoothing or underestimating low-abundance signals. Importantly, we validated the method on a benchmark dataset containing mixtures of three species’ proteomes in defined ratios, as well as on real single-cell proteomics datasets, demonstrating its robustness and broad applicability. SoftHybrid has been implemented as an R package with detailed documentation and workflow examples, and is freely available on GitHub at https://github.com/YixinShiProteomics/softHybridImpute.

## Materials and Methods

### Dataset design for benchmarking

#### Three-species mix dataset (Dataset HYE)

comprises a dilution series at 50 pg, 100 pg, 1 ng, and 10 ng, with two mixtures (Sample A vs Sample B) differing in species composition (Sample A consisted of 65% (w/w) human, 30% (w/w) yeast, and 5% (w/w) *E. coli* proteins, whereas Sample B contained 65% (w/w) human, 15% (w/w) yeast, and 20% (w/w) *E. coli* proteins; Figure 1A) followed by label-free quantitation of eight technical replicates. Receiver Operating Characteristic (ROC) curves were separately analysed across different doses and species, and the macro-ROC was defined as the average of species-specific TPRs at matched FPR across up, down, and neutral proteins at each dose. For each dose and imputation method, differential expression analysis was performed between sample A and sample B using the limma package. The resulting test statistics (t-values) were used as predictors to generate ROC curves, with yeast proteins expected to be up-regulated by 2-fold in sample B relative to sample A treated as true positives, and E. coli proteins predicted to be down-regulated by 4-fold treated as true negatives. Human proteins that were not expected to change were included as neutral controls to assess the specificity of classification.

**Figure 1.**
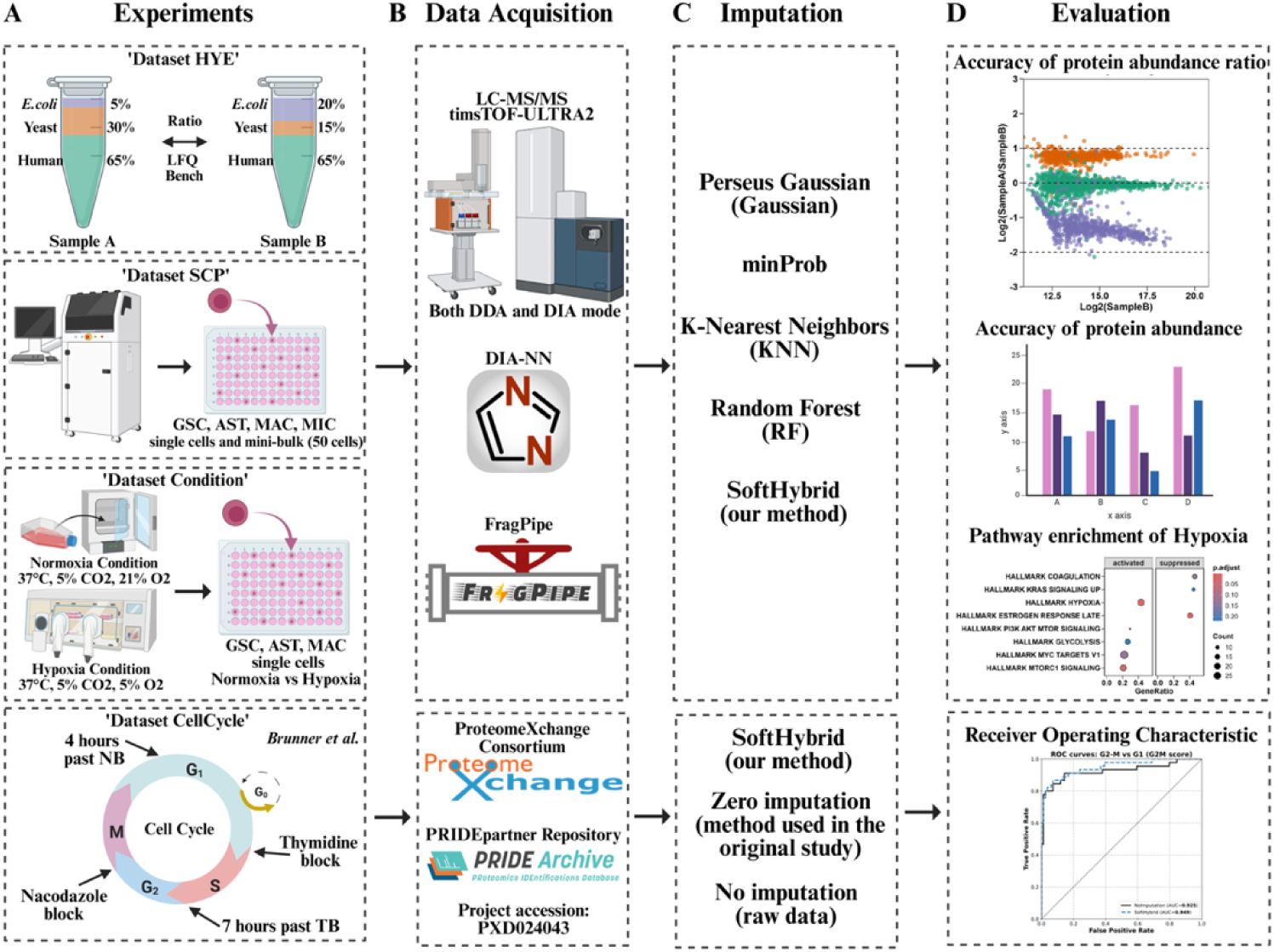
Overview of the benchmarking workflow. (A) Schematic overview of the datasets used in this study, including three in-house datasets (*Dataset HYE*: hybrid-species mixtures; *Dataset SCP*: single-cell proteomics*; Dataset Condition*: hypoxia versus normoxia conditions) and a public dataset^30^ (*Dataset CellCycle*: single HeLa cells synchronised at different cell cycle stages). (B) LC-MS/MS acquisition followed by DIA-NN or FragPipe data processing. The public dataset was retrieved from the ProteomeXchange Consortium via the PRIDEpartner repository under accession PXD024043. (C) Imputation methods benchmarked in this study (Gaussian, minProb, KNN, RF, and SoftHybrid). For the external dataset, SoftHybrid was further compared with zero-imputation (as used in the original study) and no-imputation (raw data). (D) Evaluation metrics, including abundance ratio accuracy, protein abundance recovery, pathway-level biological consistency, and cell group discrimination (ROC). Corresponding results are presented in Figures 4-7. Created with BioRender.com.

#### Baseline 4-cell-type single-cell and 50-cell mini-bulk dataset (Dataset SCP)

contains single cell proteomes from four different cell types: glioblastoma stem cell (GSC), astrocyte-like cells (AST), macrophage-like cells (MAC) and microglia-like cells (MIC), with the latter three cell types derived from induced pluripotent stem cells (iPSCs). Mini-bulk samples consist of four 50-cell aggregates per type. The 50-cell bulk proteomics profiles for each cell type served as the reference ground truth for evaluating the imputed proteome profiles in single-cell proteomics data.

#### Hypoxia vs. normoxia single-cell and 50-cell bulk dataset (Dataset Condition)

contains single-cell proteomes from three cell types (GSC, AST, MAC) from different cultivation conditions (hypoxia vs normoxia). Differential analysis between hypoxia and normoxia groups was performed using the *limma* package. All proteins were ranked by logFC values, and pathway enrichment was assessed using GSEA and the molecular signatures database (MSigDB).

#### Public dataset from a study conducted by Brunner et al. (Dataset CellCycle)^30^

contains single-cell proteomes from single HeLa cells synchronised into different cell cycle stages, including G1, G1/2, G2 and G2/M, which were obtained by timsTOF-SCP (Bruker) using DIA mode and processed using DIA-NN 1.8. The DIA-NN output results were obtained from PRIDEpartner Repository (PRIDE project accession: PXD024043).

### SoftHybrid: combining MNAR- and MAR-oriented imputations

Sigmoid weighting functions were defined using data-driven centring points (*r_o_*, *x*_0_), identified from the LOESS-fitted missingness-abundance relationship by a farthest-point detection procedure. These elbow points were used as smooth transition anchors in the weighting scheme rather than as hard thresholds:

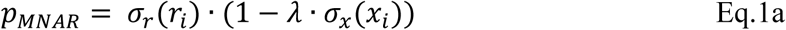

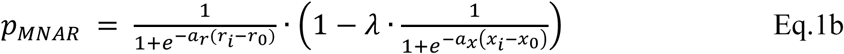

Here *a_r_* and *a_x_* set the steepness of the sigmoid weight functions and *λ* ∈ [0, 1] scales the abundance contribution. In our implementation presented herein, both *a_r_* and *a_x_* were set to 10 and 5, and *λ* was set to 0.4, providing a balanced integration of missing rate and abundance. Hyperparameter optimisation is presented in the Results).

The final imputed value for protein *i* in the sample *j* was obtained as a weighted combination of Random Forest (RF) (MAR-oriented) and minProb (MNAR-oriented):

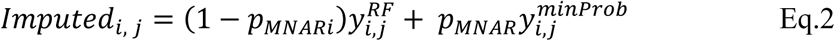

In Eq.2, 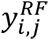 denotes the imputed value of protein *i* in sample *j* obtained using the RF method, whereas 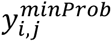 represents the corresponding value imputed with the minProb algorithm. In Supplementary Figure 1, we summarise the full processing workflow, from QC, filtering, transformation, and centring through imputation and downstream statistics. This shows how SoftHybrid integrates into both single-cell and bulk proteomics pipelines, establishing the rationale for SoftHybrid and its practical implementation.

### Cell culture

#### E57 glioblastoma stem cell culture

E57 mesenchymal glioblastoma stem cells were sourced from the Glioma Cellular Genetic Resource in Edinburgh^31^. E57 cells were cultured in DMEM/F12 (Thermo Fisher Scientific, 31330-038) supplemented with 1.5g/L glucose (Merck, G8644-100ML), MEM NEAA 1x final (Life Technologies, 11140-035), B-27 supplement 0.5x final (Thermo Fisher Scientific, 17504044), N-2 supplement 0.5x final (Thermo Fisher Scientific, 17502048), 100µM 2-Mercaptoethanol (Thermo Fisher Scientific, 31350010), 0.01% Bovine Albumin Fraction V (Thermo Fisher Scientific, 15260037), 10ng/ml Recombinant Murine EGF (PeproTech, 315-09), 10ng/ml Recombinant Human FGF (PeproTech, 100-18C), and 2µg/ml Laminin (Bio-Techne 3446-005-01). Cells were cultured in U-Shaped T75 Flasks (Corning, 430641U), coated with 10 μg/mL laminin diluted in PBS-Ca/Mg (Gibco, 15374875). Culture conditions were maintained at 37°C with 5% CO2 in a humidified CellXpert® incubator.

#### iPSC cell culture

KOLF2.1S iPSC cells were a gift from Dr Sally Cowley (Head, James and Lillian Martin Centre for Stem Cell Research University of Oxford). iPSCs were cultured in OxE8 medium^32^ on Geltrex (Life Technologies, A1413302) coated tissue culture plates. iPSCs were fed daily and passaged when reached 80% confluency using 0.5mM EDTA.

#### Macrophage,/microglia differentiation from iPSC cells

KOLF2.1S iPSCs were differentiated into macrophage/microglia precursors as described previously^33,34^. In brief, 4 × 10^6^ cells/well were seeded into a AggreWellTM800 plate (Stemcell Technologies 34811) in embryoid body (EB) medium (OxE8 supplemented with 50 ng/mL BMP4 (PeproTech, PHC9534), 50 ng/mL VEGF (PeproTech, PHC9394), and 20 ng/mL SCF (Miltenyi Biotec, 130-096-695)) with 10µM ROCK inhibitor (Abcam, Ab120129). EBs were fed daily by completing a 75% media change with EB medium without ROCK inhibitor. On day 5, the EBs from each AggreWell were divided equally between two T75 flasks and cultured in ‘Factory’ media (XVIVO (Lonza, BE02-060F) supplemented with 1x GlutaMax (Thermo Fisher Scientific, 35050-061), 2-mercaptoethanol, Penicillin -Streptomycin(Thermo Fisher Scientific, 15140-122), 100ng/ml M-CSF (Thermo Fisher Scientific, PHC9501), and 25ng/ml IL-3 (Thermo Fisher Scientific, PHC0033)). The Factories were topped up with 10ml culture media weekly, and the floating myeloid precursor cells were harvested once a week from week 5 to week 9 by removing 50% of the culture media and replacing with equal volume fresh culture media. The precursor cells were seeded into 6-well plates at a density of 8×10^5^ cells/well and differentiated either for 1 week in macrophage medium (XVIVO supplemented with 1x Glutamax and 100ng/ml M-CSF) or for 2 weeks in microglia medium (Advanced DMEM/F-12 (Thermo Fisher Scientific, 12634-010), 1x Glutamax, 100ng/ml IL-34 (PeproRTech, 200-34), 50ng/ml TGF-β1 (PeproTech, 100-21C), 25ng/ml M-CSF, and 10ng/ml GM-CSF (Thermo Fisher Scientigic, PHC2013)). A 50% media change was performed every 3-4 days.

#### Astrocyte differentiation from iPSC cells

KOLF2.1S iPSCs were first differentiated into neural precursor cells then astrocytes as described step-by-step in STAR protocols by Perriott et al^35^.

### Conditional cultivation of E57 cells and differentiated astrocytes and macrophages

After full differentiation, GSCs, differentiated ASTs and MACs were cultured in parallel for three days under normoxic conditions (37 °C, 5% CO₂, ∼21% O₂) and in a hypoxia chamber (37 °C, 5% CO₂, 5% O₂).

### Cell harvest and preparation

200,000 cells of each type under each cultivation condition were collected into individual 1.5 mL tubes. After three washes with 1× PBS (Merck, D8537), each cell pellet was resuspended in 1 mL of degassed 1× PBS, resulting in a final concentration of approximately 200 cells/µL. The suspensions were kept on ice until single-cell aliquoting.

### Cell sorting and sample preparation

Single cells were isolated and aliquoted using a CellenONE instrument and proteoCHIP EVO 96 nanowell plates (Cellenion). A master mix was prepared by combining 120 µL HPLC-grade water (Merck, 270733), 20 µL 1 M triethylammonium bicarbonate (TEAB) buffer (Merck, 18597), 40 µL 1% n-Dodecyl-β-D-maltoside (DDM; Merck, D4641), and 20 µL 100 ng/µL MS-grade trypsin (Fisher Scientific, 13454189). One microliter of master mix was dispensed into each well, followed by single-cell dispensing. The proteoCHIP plates were incubated at 50 °C for 1 h, cooled to 20 °C, and diluted with 3 µL 0.1% formic acid (FA; Merck, 5330020050; solvent A). Peptides were transferred by centrifugation at 800 × g for 1 min into a low-binding 96-well plate (Eppendorf, EP0030129580), after which 17 µL solvent A was added to each well. Plates were sealed with foil and stored at −20 °C until Evotip loading.

### Three-species mix

Three-species proteomic mixtures were prepared following the protocol described by Navarro et al^36^. Sample A consisted of 65% (w/w) human, 30% (w/w) yeast, and 5% (w/w) *E. coli* proteins, whereas Sample B contained 65% (w/w) human, 15% (w/w) yeast, and 20% (w/w) *E. coli* proteins. Aliquots of Samples A and B were prepared at four total protein input levels (50 pg, 100 pg, 1 ng, and 10 ng). For each input level, paired Samples A and B were analysed with eight technical replicates, yielding a total of 64 samples.

### LC-MS/MS analysis

Evotips were prepared according to the manufacturer’s instructions. Samples were analysed on a timsTOF Ultra2 mass spectrometer (Bruker Daltonics GmbH) coupled to an Evosep One system. Peptide separation was performed using the Whisper 40SPD Evosep method with an integrated emitter column (IonOpticks Aurora Elite Gen 3; 150 mm × 75 µm, 1.7 µm particle size, 120 Å pore size). Full MS scans were acquired over an m/z range of 100–1700 with a 1/k₀ range of 0.64–1.37 V·s/cm², using an accumulation and ramp time of 100 ms. For all three datasets, data were acquired in dia-PASEF mode, employing 25 m/z isolation windows spanning 400–1000 m/z, resulting in a total of 8 dia-PASEF frames with 3 ion mobility windows. For *Dataset HYE*, parallel acquisitions were also performed in DDA mode, using a cycle time of 1.89 s, a target intensity of 10,000, and an intensity threshold of 1,500, with other parameters set to default values.

### Raw data processing

Raw diaPASEF data were processed using DIA-NN (v2.2.0)^37^. Protein identification was performed against the UniProt human proteome (retrieved on February 25, 2026; 20,391 entries) and a combined UniProt FASTA database containing *Homo sapiens, Saccharomyces cerevisiae, and Escherichia coli* proteomes (retrieved on February 24, 2026; 30,835 entries). Both FASTA files were reviewed and supplemented with common contaminant proteins from the common Repository of Adventitious Proteins (cRAP) database maintained by the Global Proteome Machine (GPM) resource^38^. Both library-free searching and library generation were employed, with match-between-runs (MBR) enabled. Trypsin/P was selected as the protease, allowing up to one missed cleavage. N-terminal methionine excision was included as a modification for *Dataset SCP* and *Dataset Conditio*n while both N-terminal methionine excision and C-carbamidomethylation were specified as modifications for *Dataset HYE*, with a false discovery rate (FDR) set at 1%. The peptide length was restricted to 7-30 amino acids. Precursor charge states ranged from 1 to 4, with an m/z range of 300-1800. Fragment ion m/z range was set to 200-1800. Both mass accuracy and MS1 accuracy were set to 15 ppm, with the scan window set to zero. Protein quantification was performed using the DIA-NN QuantUMS algorithm^39^, which summarises precursor-level signals to protein-level intensities with cross-run normalisation enabled by default. Data from different batches and loading amounts were processed separately.

Raw data acquired in DDA mode were analysed using FragPipe (v23.1) with the integrated MSFragger search engine and Philosopher toolkit^40,41^. Spectra were searched against the combined UniProt FASTA database (human, yeast and *E.coli*), supplemented with common contaminant proteins and decoy sequences for FDR estimation. Database searching was performed using MSFragger (v4.3) with tryptic specificity, allowing up to two missed cleavages. The precursor mass tolerance was set to ±20 ppm, and fragment mass tolerance to 20 ppm. Carbamidomethylation of cysteine residues was specified as a fixed modification, while methionine oxidation and protein N-terminal acetylation were set as variable modifications. Peptide-spectrum matches (PSMs) were validated using Percolator (v3.7.1), and protein inference was performed using ProteinProphet within the Philosopher toolkit (v5.1.2). Label-free quantification was performed using IonQuant (v1.11.11) with MBR enabled. Peptide and protein identifications were filtered to an FDR of 1%.

### Bioinformatics analysis

All bioinformatic analyses were performed in R (version 4.3.2) using RStudio. The data processing pipeline included protein filtering, log_2_ transformation, median centring, missing-value imputation, and downstream analyses such as principal component analysis (PCA), differential expression analysis, pathway enrichment and receiver operation characteristic (ROC) (Figure 1, Figure S1). Samples with fewer than 1,000 proteins identified were excluded. Proteins classified as contaminants were removed from all datasets, and proteins detected in fewer than 25% of replicates across all groups were also removed. In total, four commonly used imputation methods, one previously developed hybrid strategy, and the imputation approach in the original study were benchmarked against SoftHybrid. These include Gaussian (downshift 1.8, width/scale 0.3), minProb (*imputeLCMD* package in R with default settings, q = 0.01, tune.sigma = 1)^42^, KNN (k=10, *impute.knn* package in R)^10,43^, random forest (RF, *missForest* package in R)^16^, MSImpute (*msImpute* package in R with ‘v2-mnar’ method selected for imputation)^24^ and zero imputation. To ensure a fair and reproducible comparison, all baseline methods were implemented using either their widely adopted standard settings or default parameters, as commonly used in proteomics and omics benchmarking studies.

### Hyperparameter Optimisation and Generalisation

The hyperparameters *a_r_*, *a_x_* and *λ* were optimised using the *Dataset HYE* under both DDA and DIA acquisition modes. The search grids were set to *a_r_* ∈ {2, 5, 6, 7, 8, 9, 10, 11, 12, 13, 14, 15, 20}, *a_x_* ∈ {1, 2, 3, 4, 5, 6, 7, 8, 9, 10} and *λ* ∈ {0, 0.1, 0.2,… 1}. Performance was evaluated using the root mean square error (RMSE) between imputed log_2_(SampleA/SampleB) ratios and the corresponding ground truth ratios (GT_ratio_; 1 for yeast proteins, 0 for human proteins, and −2 for *E.coli* proteins; Eq.3). To assess hyperparameter generalisability, the external public *Dataset CellCycle* was analysed using the same grid search strategy. Performance was evaluated by the area under the receiver operating characteristic (ROC) curve (AUC) of G2M marker-based scoring for distinguishing G2M from G1 cells.

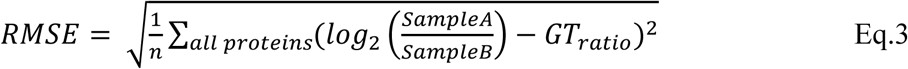

To visualise the effects of individual hyperparameters on imputation performance, grid search results and RF weights were evaluated using *Dataset SCP* with masked 50-cell data. Masking was performed following a previously reported benchmarking strategy^8^. Hyperparameter combinations of *a_r_* ∈ {2,8}, *a_x_* ∈ {2,8} and *λ* ∈ {0.2, 0.8} were applied to illustrate their influence on the resulting weighting profiles.

### Benchmarking Analysis

In *Dataset HYE*, imputation methods were benchmarked based on absolute error (AE, Eq.4) and mean absolute error (MAE, Eq.5) between imputed ratios and ground truth, ratio scatter plots, and species separation performance assessed by ROC analysis, under both DDA and DIA acquisition modes. Statistical significance was evaluated using the Wilcoxon test for AE comparisons and the DeLong test for AUC comparisons. P-values were adjusted using the Benjamini-Hochberg method.

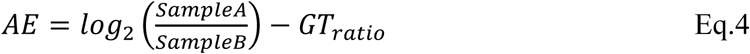

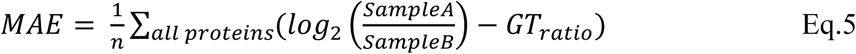

In *Dataset SCP*, cell-type-specific proteins (Class A proteins) were defined as proteins with a detection rate of 100% in at least one cell type and a missing rate of 100% in one another cell type within the 50-cell mini-bulk dataset. Fold change (FC) was defined as the difference between the mean protein abundance among expressing cell types and that in non-expressing cell types. To evaluate imputation performance for these proteins, the Spearman correlation between FC values derived from imputed single-cell data and the mean protein abundance in expressing cell types within 50-cell mini-bulk dataset was assessed. Spearman correlation between the imputed single-cell matrix and the original raw data matrix was used to evaluate overall similarity to the original data after imputation. Pearson correlation among replicates of the same cell type was calculated to assess the extent of variability introduced by imputation. In addition, clustering performance and preservation of cell-cell geometry were evaluated using Adjusted Rand Index (ARI)^44^, Normalised Mutual Information (NMI)^45^, and Mantel test^46^.

In *Dataset Condition*, pathway enrichment analysis was performed to evaluate the recovery of the target HALLMARK HYPOXIA pathway across different imputation methods. Gene set enrichment analysis (GSEA) was conducted using genes sets from the Molecular Signatures Database (MSigDB)^47^. Enrichment was evaluated based on the normalized enrichment score (NES) and gene ratio derived from the GSEA results. P-values were adjusted using the Benjamini-Hochberg method. Hypoxia proteins were defined as the proteins annotated in the HALLMARK HYPOXIA gene set and detected in *Dataset Condition*. The mean abundance shift was calculated as the difference in mean protein abundance between normoxia and hypoxia conditions.

Generalisation analysis was conducted on the public *Dataset CellCycle*. Receiver operating characteristic analysis was conducted using cell-cycle-specific markers to distinguish between different cell cycles in the SoftHybrid imputed dataset, and the results were compared to the originally used one (zero imputation) and the non-imputed raw dataset, using the DeLong significant test. The cell-cycle-specific marker list was obtained from the original study’s supplementary materials (https://github.com/theislab/singlecell_proteomics)^30^.

## Results

### Dataset preparation for benchmarking

We assembled three datasets (*Dataset HYE, Dataset SCP and Dataset Condition*) to benchmark imputation in complementary settings: a controlled three-species titration (*Dataset HYE*) to provide known fold-change ground truth; a four-cell-type single-cell panel *(Dataset SCP*) to test clustering fidelity and replicate consistency under high sparsity; and a single-cell perturbation (*Dataset Condition*: hypoxia vs. normoxia) to assess biological interpretability. For Dataset HYE, we acquired data in both DIA and DDA modes to probe method robustness across distinct missingness regimes (DIA typically has a lower missing rate, while DDA has a higher missing rate with more MAR-like characteristics due to dynamic exclusion/precursor competition). Global and per-sample missing rates increased at lower input and were consistently higher in DDA than DIA (Figure S2). An external public dataset *(Dataset CellCycle*) was additionally included for method optimisation and to assess its broader applicability.

#### Dataset HYE

The expected log_2_ (A/B) species-level fold changes are –2 (*E. coli*), +1 (yeast), and 0 (human). For each input level, we acquired 16 replicates split evenly between DIA (n=8) and DDA (n=8).

#### Dataset SCP

It contains 20 single cells per type plus four 50-cell aggregates per type. In total, 2,876 proteins were detected across the single-cell set and 7,375 proteins across the 50-cell aggregates. After standard filtering and quality control (Figure S1), 76 single cells were retained for downstream analysis, and all 50-cell samples were retained.

#### Dataset Condition

It profiles three cell types under hypoxia and normoxia, with 20 single cells per condition per type (total intended n=120). We detected 3,842 proteins across the experiment; 106 cells passed quality control and entered downstream analysis.

#### Public Dataset CellCycle^30^

It contains HeLa single cells synchronised to different cell cycle stages (G1, G1/S, G2, G2/M) and after filtration, 424 single cells and 1867 proteins were maintained for downstream analysis.

### Imputation choices have effects on downstream biological interpretation

We first assessed how different imputation strategies influence downstream proteomics analysis in *Dataset SCP*. Different imputation methods were applied to the same data (*Dataset SCP*), and the resulting imputed matrices were combined for PCA visualisation. The PCA revealed that the primary source of variation (PC1) was driven by differences between imputation strategies (Figure 2A). To further investigate this observation, we examined the relationship between absolute PC1 loadings and protein-level missing rates and found a strong positive correlation (*r* = 0.957, *p* < 0.001, Figure S3A), indicating that proteins with higher missingness contribute more to the separation along PC1. Although the overall cell-type structure remains comparable across imputation methods (Figure S3B-C), the dominant axis of variation is largely driven by imputation-specific effects. As shown in Figure S3D, PCA separation becomes increasingly driven by the imputation method as protein-level missingness increases. This indicates that, in sparse single-cell proteomics data with a high missing rate, method-specific imputation behaviour can strongly influence the variance structure of the completed data matrix. Taken together, these findings suggest that different imputation strategies can introduce systematic biases into the data, leading to divergent representations of the same dataset in downstream analysis, highlighting that imputation should not be considered as a neutral pre-processing step^8,10^.

**Figure 2.**
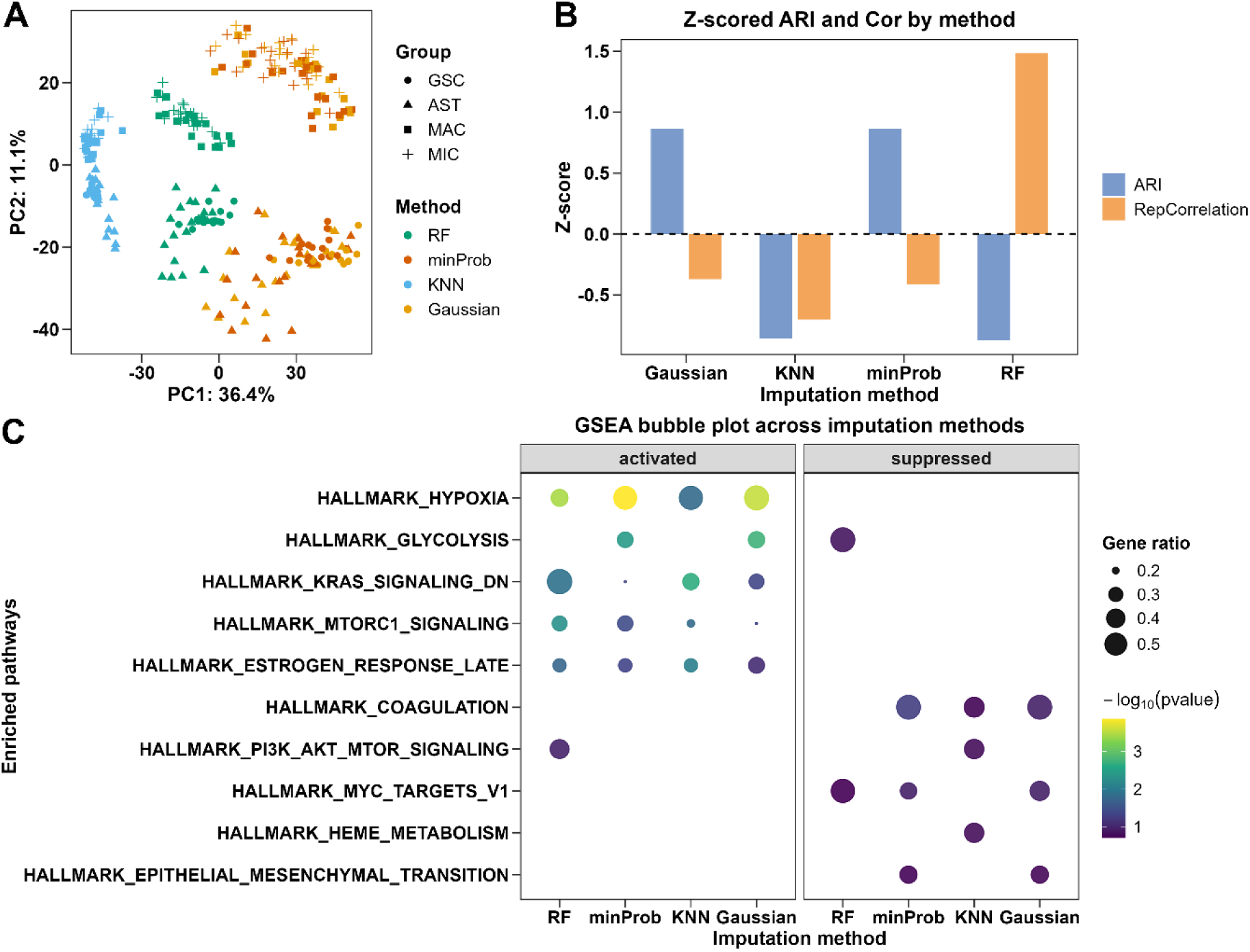
Imputation methods are not neutral procedures. (A) PCA of proteomics data imputed with commonly used methods (RF, minProb, KNN, and Gaussian) across four cell types (GSC, AST, MAC, and MIC) using the same single-cell proteomics dataset (*Dataset SCP*; Separated PCA plots per imputation method was shown in Figure S3C). (B) Quantitative comparison of imputation methods based on ARI and replicate Pearson correlation, shown as Z-scores. (C) GSEA results of limma differential analysis, illustrating enriched hallmark pathways that are either activated (left) or suppressed (right). Dot size represents gene ratio, and colour indicates statistical significance (-*log*_10_ p-value).

To quantify this effect, we compared clustering fidelity and replicate consistency using the Adjusted Rand Index (ARI) and replicate Pearson correlations (Figure 2B). RF yielded the highest reproducibility across replicates, but compressed real biological heterogeneity between cell types, obscuring group-level differences with low ARI. KNN over-smoothed the data, leading to poor clustering agreement. However, Gaussian and minProb better preserved between-group variation at the cost of reduced replicate concordance and cell-cell geometry preservation (Figure 2B, Figure S3B). In short, none of these widely used methods simultaneously achieve robust separation of distinct cell populations and strong within-group agreement, both essential for single-cell annotation and inter-group comparisons.

We next examined the functional consequences of different imputation strategies in *Dataset Condition* by contrasting normoxic and hypoxic conditions using GSEA (Figure 2C). As expected, hallmark pathways directly related to hypoxia, such as *HALLMARK HYPOXIA* were consistently recovered across all methods; however, detection sensitivity varied, reflecting the inconsistent capture of true hypoxia responses. minProb yielded the highest statistical significance for *HALLMARK HYPOXIA,* and Gaussian retrieved the largest number of hypoxia-related proteins, suggesting a greater sensitivity in recovering true biological changes. Yet this heightened sensitivity may also reduce specificity, increasing the likelihood of false positives. In line with this, pathways known to interact with hypoxia responses, such as *HALLMARK GLYCOLYSIS, HALLMARK MTORC1 SIGNALING, HALLMARK PI3K AKT MTOR SIGNALING,* and *HALLMARK MYC TARGETS,* were differentially enriched depending on the imputation strategy. More concerningly, pathways with only weak or indirect links to hypoxia, including *HALLMARK COAGULATION*, *HALLMARK ESTROGEN RESPONSE LATE,* and *HALLMARK KRAS SIGNALING_DN,* were also frequently identified, raising the possibility of spurious enrichments induced by imputation artefacts. Together, these findings demonstrate that imputation choices propagate variability from clustering and replicates into pathway-level results, thereby limiting the reliability of biological insights.

In summary, existing imputation methods may not fully preserve both inter-group differences and intra-group consistency, leading to variability in pathway-level interpretations. This limitation is particularly relevant in single-cell proteomics, where accurate annotation and reliable comparisons across cell populations are critical, especially when analysing unknown or heterogeneous cell types.

To address these challenges, we developed an imputation framework that explicitly considers the mixed nature of MAR and MNAR values, with the aims of improving the robustness of downstream analyses. Based on our analyses and previous studies^8,48^, RF and minProb showed strong performance within their respective categories (Figure 2A-C); accordingly, our strategy integrates RF for imputing MAR-like values and minProb for imputing MNAR-like values.

### Development, optimisation, and generalisation of the SoftHybrid Imputation method

In *Dataset SCP*, we observed a strong association between protein-level missing rate and biological derived missingness, as well as a moderate relationship between protein abundance and biological missingness. Notably, the overall missing rate showed a stronger correlation than protein abundance (*r* =0.95 vs *r* = −0.58, Figure 3A-B). Based on these observations, missing rate was used as the primary proxy, while protein abundance was incorporated as a secondary modulating factor (Eq.1a). In addition, the relationship between missing rate and protein abundance exhibited a data-driven elbow point (Figure 3C), suggesting a potential transition in missingness behaviour around the detection limit. Motivated by this pattern, we employed a sigmoid function to approximate the relationship between these proxies and the proportion of MNAR-like values in the hybrid model (Eq.1b). The resulting weighting scheme of the SoftHybrid model (Figure 3D) showed a smooth transition between RF-dominated (MAR-like) to minProb-dominated (MNAR-like) imputation, consistent with the intended design.

**Figure 3.**
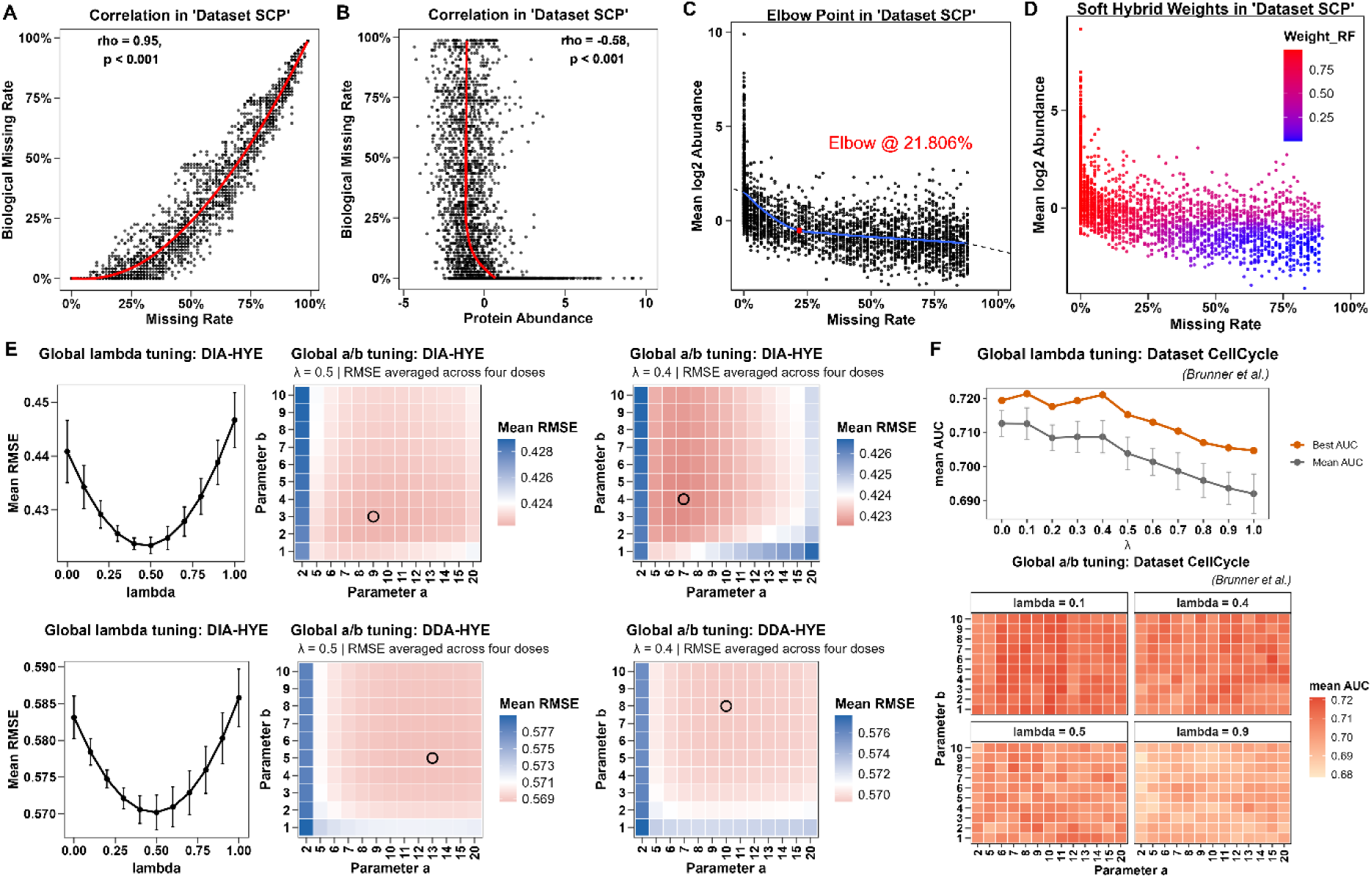
Concept, implementation, and hyperparameter optimization of the SoftHybrid imputation strategy. The SoftHybrid framework and its optimisation are illustrated using *Dataset SCP*, *Dataset HYE*, and the public *Dataset CellCycle*. (A) Predicted biological (MNAR-like) missing rate plotted against overall missing rate, showing a significantly high positive correlation with a sharp increase near an elbow point. (B) Correlation between predicted biological missing rate and protein abundance, revealing a negative relationship, with a clear transition point. (C) Identification of the missing-rate elbow point (highlighted in red, *r*_0_ = 21.806%) using LOESS regression with a maximum-distance method. (D) The SoftHybrid framework integrates RF and minProb imputations via a sigmoid-based MNAR weighting scheme. The weight distribution shows a smooth transition from RF-dominant to minProb-dominant imputation around the elbow point. (E) Hyperparameter optimisation in *Dataset HYE*. Global tuning showed that *λ* = 0.5 achieved the lowest RMSE, while the effects of *a_r_* (parameter a), and *a_x_* (parameter b) were comparatively modest. RMSE plateaued around *a_r_* ≈ 10 and *a_x_* ≈ 5. (F) Hyperparameter generalisation in the public *Dataset CellCycle*. The global mean AUC remained relatively stable for *λ* values between 0.1 and 0.4, followed by a gradual decline beyond 0.5. High-performing parameter combinations were distributed across the *a_r_*-*a_x_* space, with *a_r_* = 10 and *a_x_* = 5 yielded consistently competitive performance across different *λ* settings.

To further optimise the performance of SoftHybrid, hyperparameter grid searches were conducted in both *Dataset SCP* and *Dataset CellCycle*. The results suggest that *a_r_* primarily controls the steepness of the transition with respect to missing rate, whereas *a_x_* modulates the rate of change along protein abundance (Figure S4A-C). The parameter *λ* governs the relative weighting in the transition region, with higher values leading to increased contribution from RF (Figure S4A, D).

In *Dataset SCP*, where performance was evaluated using RMSE, *λ* = 0.5 yielded the best results, while *λ* = 0.4 provided comparable performance across both DIA and DDA modes. The optimal values for *a_r_* and *a_x_* varied between acquisition modes, with DIA favouring smaller values and DDA favouring larger values; however, a shared near-optimal region was observed around *a_r_* = 10 and *a_x_* = 5 (Figure 3E).

In contrast, in *Dataset CellCycle*, where performance was assessed using a classification-based metric, smaller values of *λ* were generally preferred, with performance declining beyond *λ* = 0.4. In comparison, *a_r_* and *a_x_* appeared to have relatively limited influence on the overall results (Figure 3F). This trend is consistent with the increased contribution of minProb at lower *λ*, which may enhance separation between cell groups by assigning lower values to missing entries. To achieve balanced performance across diverse analysis tasks in single-cell proteomics, we selected a compromise parameter set of *λ* = 0.4, *a_r_* = 10 and *a_x_* = 5 as the default configuration. The data-driven elbow point is estimated automatically within the algorithm for each input dataset.

### Benchmarking imputation strategies across single-cell-equivalent to bulk proteomics using three-species titration

To extend the evaluation of imputation performance beyond single-cell datasets, we conducted a systematic benchmarking on three-species proteomics samples across a titration series ranging from single-cell-equivalent doses (50 pg) to bulk-like levels (10 ng). As expected, the overall missing rate decreased with increasing sample amounts and was consistently higher in DDA compared with DIA (Figure S2), representing different scenarios in real-world datasets. For the single-cell-equivalent doses, the missing rates for DIA and DDA modes were 0.19 and 0.23. Further analysis revealed that missing rates at these elbow points (determined using LOESS regression with a maximum-distance method) ranged between 10% and 20%, with the precise values varying across doses and acquisition modes, reflecting experimental differences in the relative contributions of MAR and MNAR (Figure S5). Across all doses and acquisition modes, SoftHybrid adapted the balance between MAR- and MNAR-oriented imputation through the underlying sigmoid-based weighting scheme, with the weights changing smoothly around dataset-specific elbow points in missing rate and abundance, yielding a continuous transition between the two regimes (Figure S6).

Global error analysis revealed that SoftHybrid consistently achieved the lowest mean absolute error (MAE) across both DIA and DDA datasets (Figure 4A, D). In the DIA dataset, these improvements were statistically significant across all input levels, with the advantage being most pronounced at the lowest doses where missing values were most prevalent, indicating improved performance under highly sparse conditions. In contrast, in the DDA dataset, although SoftHybrid generally exhibited lower MAE, the differences were not statistically significant at the 100pg dose when compared with Gaussian and minProb, suggesting that under certain conditions, these three methods can achieve comparable overall accuracy.

**Figure 4.**
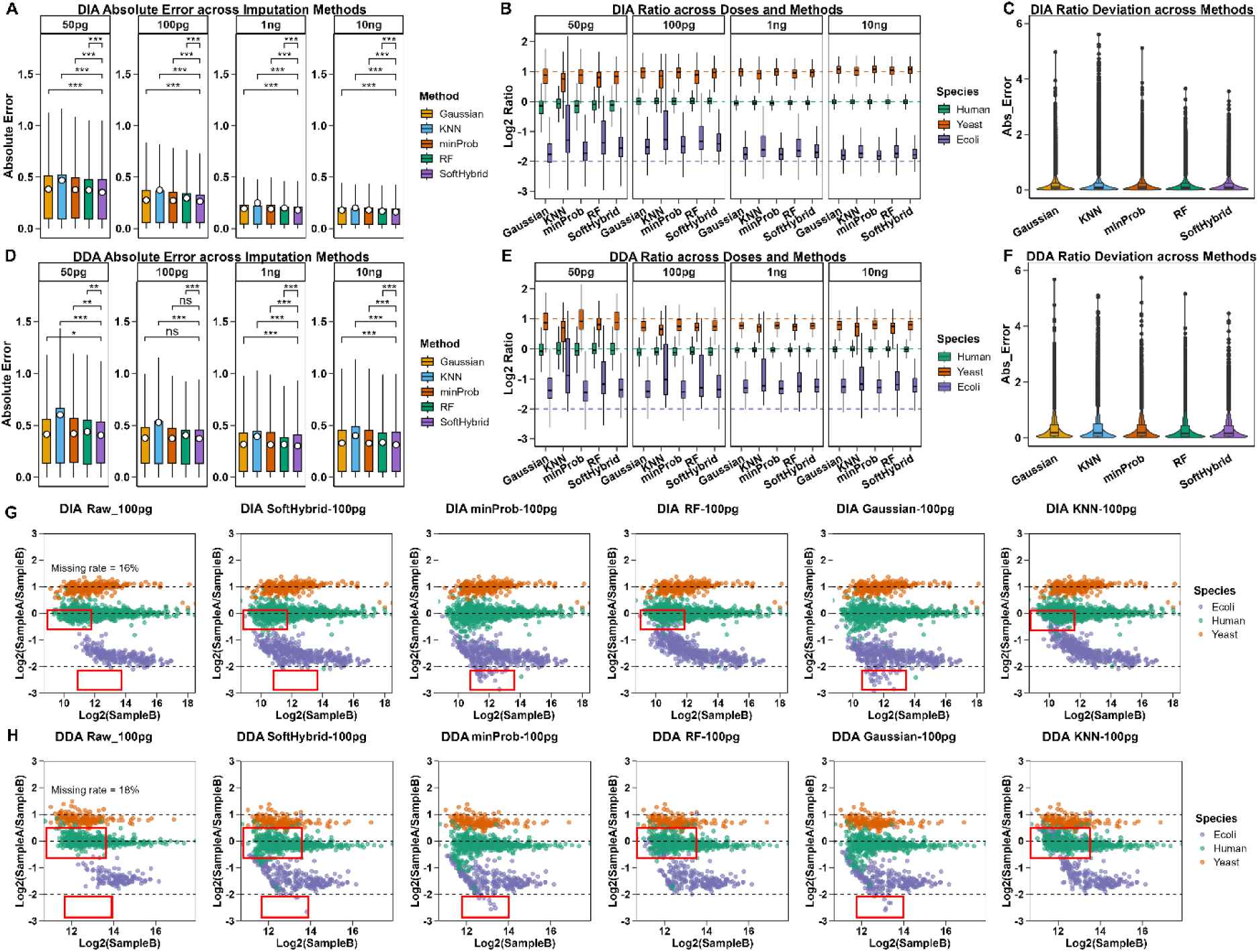
**Benchmarking of imputation methods based on AE, log2 ratio estimation and deviation from ground truth across doses and species in *Dataset HYE* under DIA and DDA acquisition modes**. (A, D) Distribution of absolute error (AE) across imputation methods at different protein input amounts (50pg, 100pg, 1ng, 10ng) for DIA (A) and DDA (D) datasets. Boxes represent interquartile ranges, with white circle indicating means; whiskers denote data range. Statistical significance was assessed using the Wilcoxon test (***, *p* < 0.001; **, *p* < 0.01; *, *p* < 0.05; ns, *p* > 0.05). (B, E) Species-wise log2 ratio estimates across imputation methods at each input level for DIA (B) and DDA (E). (C, F) Distribution of absolute error between log2 ratios and ground truth for individual proteins. (G-H) Scatter plots of log2 ratios versus log2(Sample B intensities) at 100 pg for DIA (G) and DDA (H). Representative results are shown for raw data, SoftHybrid, minProb, RF, Gaussian, and KNN. For raw data, proteins with missing values were excluded prior to ratio calculation, resulting in substantially reduced protein set (total protein number are provided in Figure S9). Dashed horizontal lines indicate the expected ground truth *log*_2_ ratios (human: 0; yeast: 1; *E.coli*: −2). Red boxes highlight regions where systematic biases are present or corrected by different imputation methods.

To better understand whether this advantage held across different biological contexts, we next stratified the analysis by species. As expected, human proteins, which exhibited the highest protein abundance showed the most accurate log2 ratio estimates, followed by yeast, whereas *E. coli* proteins, contributing the lowest absolute protein amounts, displayed the largest deviations from the expected values, particularly at lower input levels in both DIA and DDA datasets (Figure 4B, E). At the method level, distinct behaviours were observed. RF and KNN consistently oversmoothed the imputed data, thereby leading to ratio compression across all species, most prominently for *E. coli*. In contrast, minProb and Gaussian imputation produced median log2 ratios closer to the expected values. However, these methods also showed increased variability at the protein level, as reflected by the more pronounced extreme deviations (Figure 4C, F). SoftHybrid achieved near-optimal performance in log2 ratio estimation while maintaining the lowest overall deviation. Notably, minProb and Gaussian introduced systematic underestimation at intermediate abundance levels, whereas RF and KNN tended to overestimate in low-abundance regions, SoftHybrid mitigated these biases across the full abundance spectrum (Figure 4G-H; Figure S7-8; highlighted by red boxes). Taken together, the results indicate that SoftHybrid achieves a more balanced trade-off between bias and variance by limiting extreme deviations while maintaining reasonable agreement with expected ratios. As a result, SoftHybrid achieves improved or comparable accuracy across datasets and input levels, with the greatest benefit observed under conditions of higher data sparsity.

To further evaluate detection sensitivity and differential expression accuracy, we constructed macro-averaged ROC curves using yeast as the positive class, *E. coli* as the negative class, and human proteins as neutral controls. In DIA mode, SoftHybrid consistently delivered the highest AUC values at the lowest input levels (50 pg and 100 pg), where it clearly outperformed conventional approaches, while at higher doses (1 ng and 10 ng) its performance remained comparable to that of RF (Figure 5A; Figure S10A). In DDA mode, SoftHybrid likewise showed strong and stable performance across all tested amounts, yielding the highest mean macro-AUC and maintaining robustness throughout the dilution series (Figure 5B; Figure S10B). Across individual classification tasks, SoftHybrid either significantly outperformed or showed comparable performance relative to the other four methods (Figure S10A-B; Table S1-2). A small number of exceptions were observed. At higher input levels (1ng and 10ng) in DIA mode, RF achieved significantly higher performance for human proteins, whereas at 100pg in DIA mode, Gaussian performed better for *E.coli*. These differences are consistent with differences in missingness mechanisms across conditions. At higher doses, missing values in human proteins are more likely to resemble MAR, where typically RF-based approaches perform well. In contrast, for low-abundance *E.coli* proteins at lower input levels, missingness is more consistent with MNAR, favouring lower-tail imputation strategies such as Gaussian and minProb.

**Figure 5.**
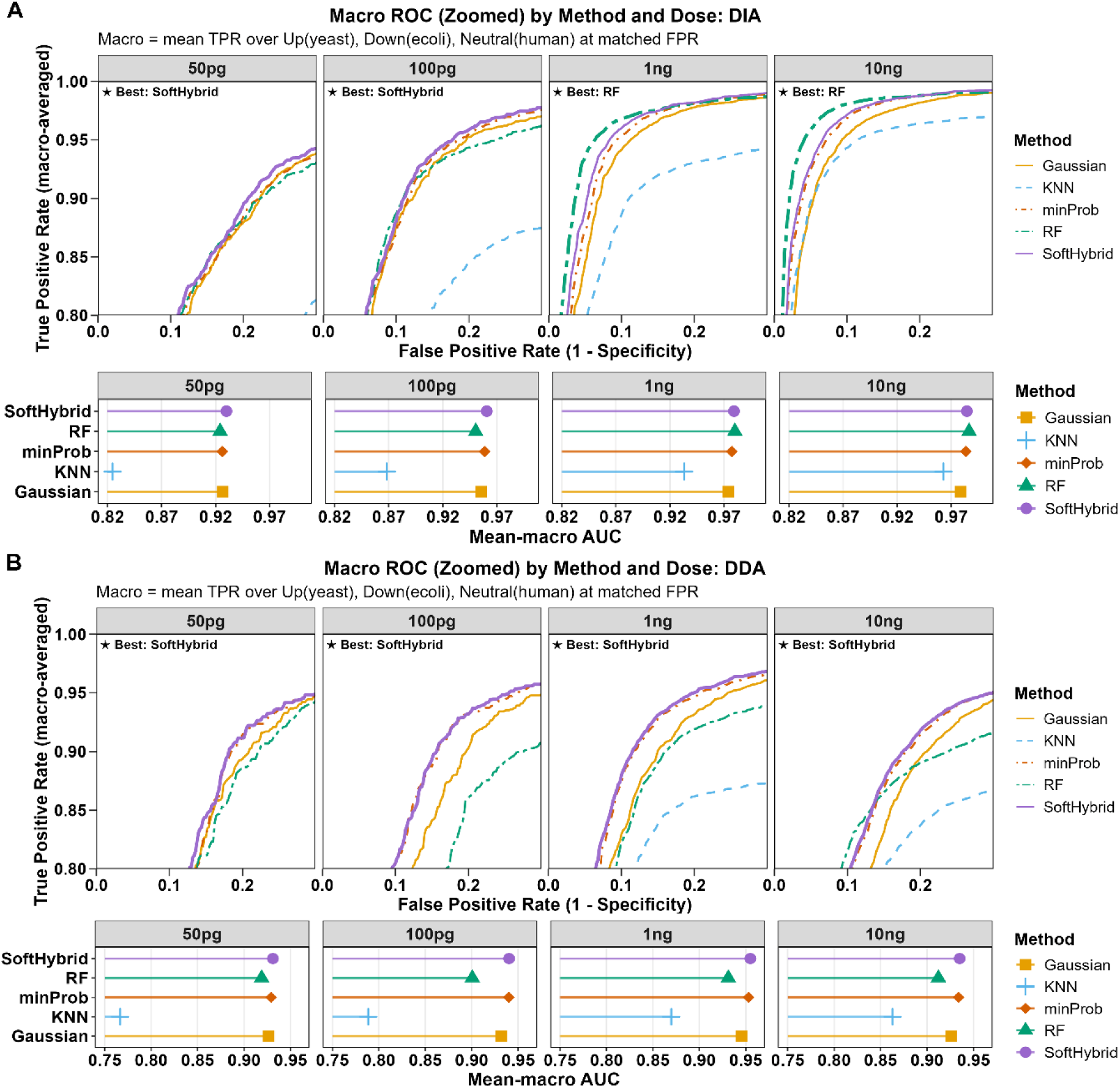
Macro-ROC analysis of imputation methods across doses in the three-species dataset. (A–B) Macro-ROC curves for four protein input amounts (50pg, 100pg, 1ng, 10ng) under DIA (A) and DDA (B) acquisition modes. The macro true positive rate (TPR) was calculated as the mean TPR across upregulated (yeast), downregulated (*E. coli*), and neutral (human) proteins at matched false positive rates (FPR). Insets summarise macro-AUC for each method at each dose, with the best-performing method indicated. ROC curves are shown in a zoomed view; full, unzoomed curves are provided in Figure S10. Statistical comparisons were performed using the DeLong test, with detailed results reported in Table S1-2.

Taken together, these results show the robustness of SoftHybrid across experimental scales, from single-cell proteomics to bulk-level datasets, with particularly strong performance at low-input levels. By adaptively balancing MAR and MNAR imputation, SoftHybrid reduces imputation-induced bias from inappropriate MNAR/MAR assumptions and minimises false discoveries, providing a reliable and broadly applicable solution for complex proteomics scenarios.

### Performance evaluation in real-world single-cell proteomics datasets

Having established SoftHybrid as suitable imputation methodology in a controlled ground-truth benchmark scenario (three-species mix), we next assessed its performance in real biological contexts using single-cell proteomics datasets, beginning with our *Dataset SCP*. Given the growing interest in cell-type deconvolution based on the proteome, we specifically evaluated imputation performance for such candidate cell-type-specific proteins across all methods (Figure 6A). Structure-based imputation methods (RF and KNN) markedly disrupted the relationship between imputed FC estimates and bulk mean expression within the expressing cell (*ρ* = 0.174 for RF and *ρ* = −0.126 for KNN), suggesting that imputing missing values in non-expressing cell types using information learned from expressing groups leads to an artificial attenuation of the contrast between expressing and non-expressing cell types. In contrast, Gaussian and minProb achieved higher correlations ( *ρ* = 0.56 and *ρ* = 0.576 respectively), likely because their low-value imputation strategy better preserves the expected absence signal in non-expressing cell types. However, these approaches remain suboptimal when missingness occurs within expressing cell types, as imputing low values in such cases can underestimate true expression levels. Notably, SoftHybrid achieved the highest correlation (*ρ* = 0.607), mitigating the opposing biases of the two individual methods, thereby better preserving fold-change relationships for cell-type-specific proteins.

**Figure 6.**
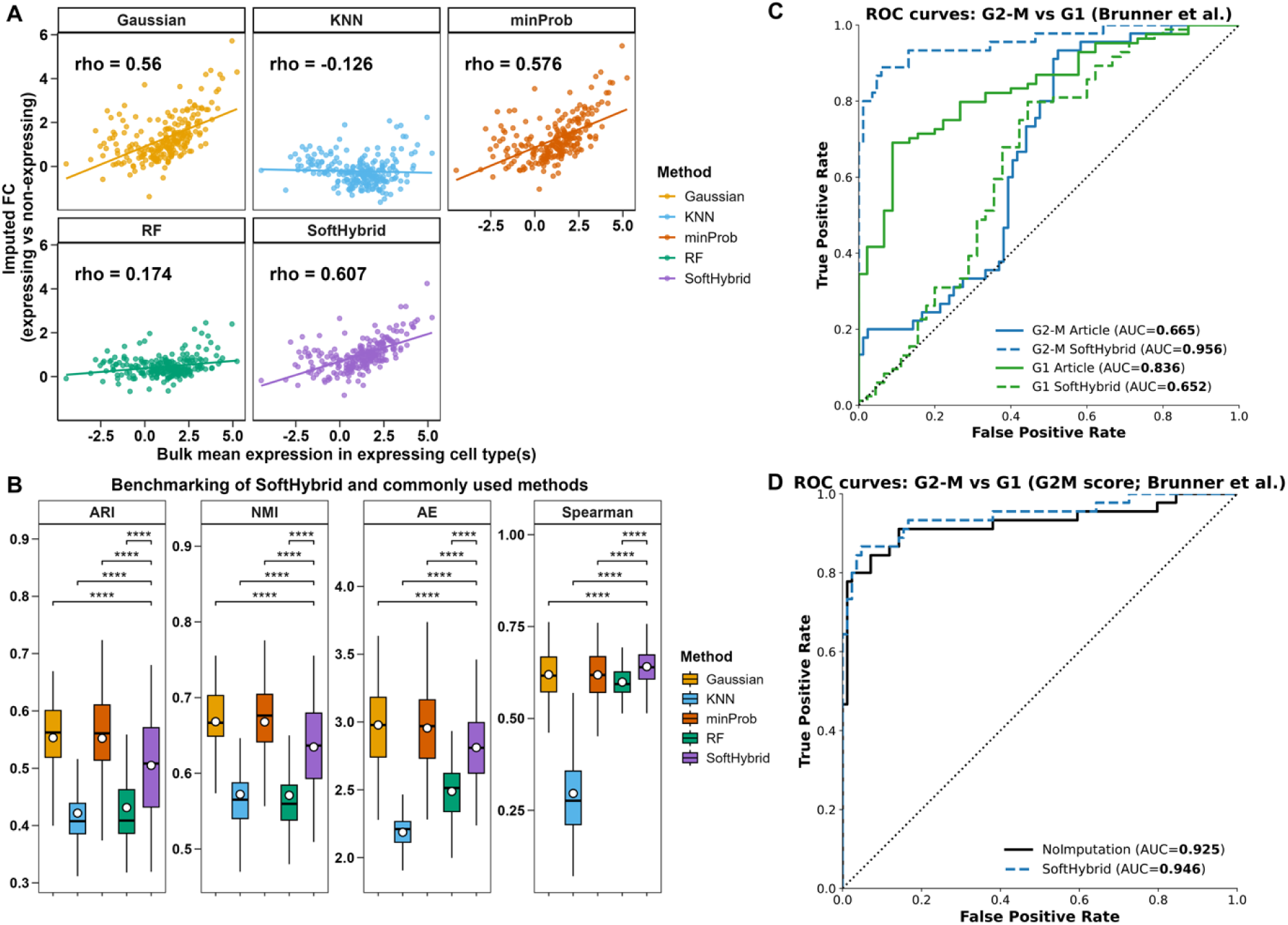
Benchmarking of imputation methods in the real-world single-cell proteomics datasets. (A) Correlation between fold changes (FC) of cell-type-specific proteins in imputed data and its corresponding bulk mean expression in expressing cell types across different imputation methods in *Dataset SCP*. The Spearman correlation coefficient *ρ* (*rho*) was calculated, with higher values indicating better agreement. (B) Performance evaluation of imputation methods using ARI, NMI, AE and Spearman correlation against 50-cell bulk ground truth in *Dataset SCP*. Higher ARI, NMI and Spearman correlation, and lower AE indicate better performance. (C) ROC analysis comparing SoftHybrid with the zero-imputation strategy used in the original study, based on G2M marker scoring to distinguish G2M and G1 cells in the public *Dataset CellCycle*. (D) ROC comparison between SoftHybrid and imputation-free (no-imputation) data in the same classification task. Statistical significance was assessed using the Wilcoxon test for ARI, NMI, AE and Spearman correlation, and the DeLong test for AUC comparisons (ns, *p* >0.05; *, *p* < 0.05; ***, *p* < 0.001; ****, *p* < 0.0001).

Quantitative benchmarking was further performed to assess the overall performance across different imputation methods (Figure 6B). Gaussian and minProb achieved significantly higher ARI and NMI values, likely reflecting the tendency of low-value imputation to enhance apparent separation between cell types. However, these methods showed the poorest performance in terms of AE. RF and KNN yielded the lowest AE, as their structure-based strategies estimate missing values from observed data, thereby, better recovering MAR and abundance-related MNAR signals. However, they performed worst in ARI and NMI, indicating reduced preservation of cell-type-specific structure, consistent with benchmarking results in *Dataset HYE*. Importantly, SoftHybrid demonstrated balanced performance across metrics, remaining comparable in ARI, NMI and AE while achieving the highest spearman correlation with the ground truth. It is suggested that SoftHybrid most effectively preserves the relative ordering of protein abundances while maintaining overall quantitative accuracy.

Further, we compared SoftHybrid with MSImpute^24^, another hybrid imputation method (Figure S11A). The results showed that although MSImpute achieved slightly better ARI, NMI and Spearman correlation, SoftHybrid showed significant improvement in AE, suggesting that SoftHybrid provides competitive performance in clustering and structural preservation compared to MSImpute, while offering a clear advantage in recovering absolute protein abundance.

Building on these results, we next assessed the external application of SoftHybrid using a public single-cell proteomics dataset (*Dataset CellCycle*)^30^, in which HeLa cells were synchronised across different cell cycle stages and profiled at single-cell level. Cell cycle phases were subsequently distinguished using marker proteins defined in a prior bulk proteomics study^49^. Using G2-M marker scoring to classify G2-M versus G1 cells, SoftHybrid yielded a significantly higher AUC compared to the imputation strategy used in the original study (zero imputation) (Figure 6C; AUC = 0.956 vs 0.665; p < 0.001). When G1 marker scoring was applied, however, the original method achieved a higher AUC than SoftHybrid. This difference is likely attributable to the lower specificity G1 markers, which typically exhibit smaller effect sizes (logFC < 1) compared to G2M markers that show consistently larger fold changes (logFC > 1)^30^. Under these conditions, zero imputation may potentially exaggerate differences for G1 markers, effectively amplifying weak signals and leading to an apparent improvement in classification performance despite their limited discriminative power. Consistent trends were also observed in additional classification tasks, including G2M vs G1S and G2 vs G1 cells (Figure S11B). Softhybrid was further compared with the raw data (no imputation), showing a slight non-significant improvement in AUC (Figure 6D; AUC = 0.946 vs 0.925; *p* = 0.283). Overall, these results indicate that SoftHybrid provides more reliable classification performance than zero imputations, particularly for marker sets with strong differential expression, while retaining comparable performance to the raw data without introducing adverse effects.

Finally, we evaluated the biological interpretability of *Dataset Condition* using GSEA with hallmark gene sets (MSigDB)^47^. SoftHybrid identified HALLMARK HYPOXIA as the most strongly enriched pathway, alongside glycolysis and MTORC1 signalling (Figure 7A). Pathways less directly related to hypoxia were also detected, such as HALLMARK ESTROGEN RESPONSE LATE and HALLMARK COAGULATION; however, their significance and protein coverage were comparable to other methods and were generally not statistically significant (Figure 7A; Figure S11C). For the HALLMARK HYPOXIA signature, SoftHybrid achieved the highest normalised enrichment score (NES) (Figure 7B; Figure S11D), implying improved preservation of biological effects. To better understand how imputation influences these pathway-level results, we next examined the underlying protein-level missingness and abundance patterns. Missingness among hypoxia-associated proteins was substantial and differed between normoxia and hypoxia conditions (Figure 7C), suggesting that imputation strategies may introduce bias into the downstream biological interpretation. In line with this, SoftHybrid produced fold change estimate closest to the raw data (Δ = 0.86 vs 0.78; Figure 7D, Figure S11E). Conversely, RF and KNN attenuated these differences (Δ = 0.63 and 0.45), whereas minProb and Gaussian amplified them (Δ = 1.04 and 1.06). This consistent pattern highlights the distinct biases introduced by the commonly used imputation methods and underscores the importance of balancing these effects for more accurate downstream interpretation.

**Figure 7.**
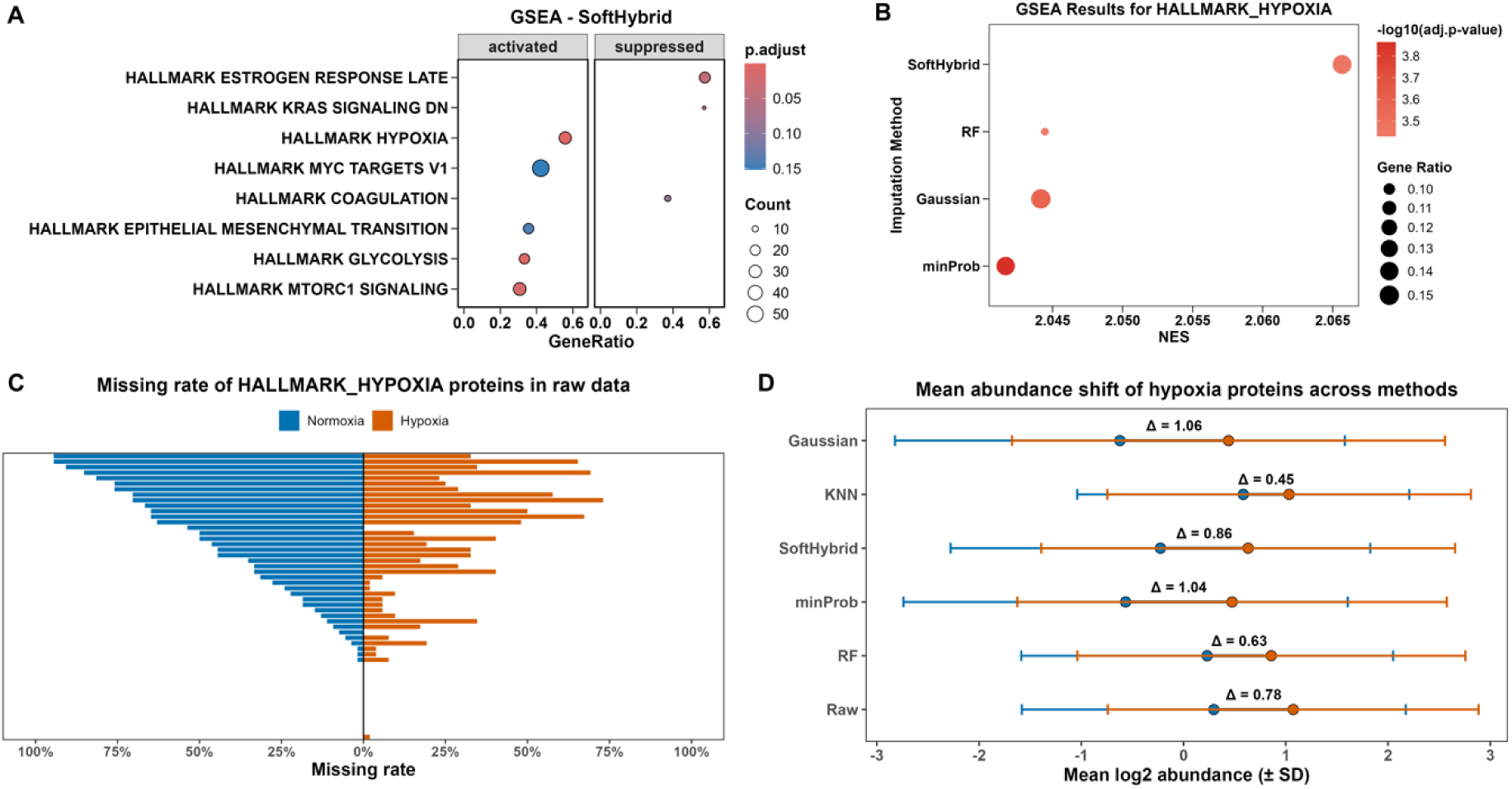
Impact of imputation methods on downstream biological interpretation. (A) GSEA of Soft Hybrid-imputed data in *Dataset Condition*. Dot size represents the number of leading-edge proteins; and colour indicates adjusted *p*-values. (B) Comparative GSEA results for the HALLMARK HYPOXIA pathway across imputation methods in *Dataset Condition*. Dot size represents gene ratio, and colour indicates −*log*_10_(adjusted *p*). (C) Missing rates of proteins associated with HALLMARK HYPOXIA pathway in raw data under normoxia and hypoxia conditions. (D) Mean *log*_2_ abundance of hypoxia-related proteins across imputation methods under normoxia and hypoxia conditions. Error bars represent standard deviation. Δ denotes the difference in mean *log*_2_ abundance between normoxia and hypoxia groups.

Taken together, these findings show that SoftHybrid not only delivers quantitative accuracy but also improves the interpretability of single-cell proteomics data, achieving these gains without inflating spurious biological signals, thereby overcoming the limitations associated with conventional imputation methods.

## Discussion

Missing values remain one of the most persistent challenges in proteomics data analysis. Because most downstream statistical tools require complete data matrices, missingness directly affects reproducibility, statistical power, and the reliability of biological interpretation^10,48^. Over the past decade, a wide range of imputation strategies have been developed to mitigate this issue, each with strengths and weaknesses. However, current methods remain unbalanced: approaches that perform well for MAR values tend to compromise biological heterogeneity, whereas those optimised for MNAR values often sacrifice replicate consistency. This problem is particularly pronounced in single-cell proteomics, where data sparsity is much more common and no systematic benchmarking of existing methods has been carried out, despite the rapid expansion of single-cell proteomics as a field. Due to the high sparsity of single-cell proteomics datasets, different imputation strategies can assign substantially different values to missing entries. When the same dataset is visualised after applying multiple imputation methods, these method-specific differences may dominate low-dimensional projections, as observed in Figure 2A. Although in practice a single imputation method is typically used for downstream analysis, differences between imputation strategies may still have effects on biological interpretation, highlighting the need for systematic benchmarking and more robust imputation approaches for single-cell proteomics.

In this study, we developed SoftHybrid, a hybrid imputation strategy that integrates missing rate and protein abundance within a continuous weighting framework. By doing so, SoftHybrid enables a smooth transition between MAR- and MNAR-oriented models, rather than relying on sharp thresholds or discrete binning. Several features distinguish this approach. First, it is applicable to both bulk and single-cell datasets (DDA and DIA), providing a unified framework across different proteomics scales. Second, SoftHybrid shows favourable performance at low-input levels and achieves comparable performance at higher-input conditions, while maintaining proteomic structure and reducing spurious signals in comparative analyses. Finally, SoftHybrid supports the recovery of biologically meaningful signals in downstream analyses, facilitating more reliable detection of pathway-level differences between biological groups.

To contextualise the design of SoftHybrid within the current landscape of hybrid imputation strategies, we compared its underlying principles with several representative approaches. The BIND strategy^22^ emphasised the biological information embedded in missing values but did not perform imputation; as a result, technical missing values may be under-accounted for, potentially compromising statistical robustness. By contrast, SoftHybrid incorporates information from both biological and technical missingness, facilitating a more balanced recovery of underlying biological signals. The bin-based strategy^23^ partitions features according to missing rate and abundance, but treats all proteins within a bin as having similar missing-value compositions. This simplification may introduce discontinuities at bin boundaries and lead to bias. In addition, the requirement for per-bin benchmarking across methods to select the optimal model may limit its transferability between datasets. However, SoftHybrid adopts a continuous weighting scheme, thereby better accommodating the mixed nature of missingness. Furthermore, as the employed weighting scheme is data driven and estimated for each dataset rather than manually defined, SoftHybrid offers unbiased and improved adaptability across diverse data structures. Consistent with this, results from external validation show that SoftHybrid can be effectively applied to independent datasets.

Among recent hybrid approaches, MSImpute^24^ was directly benchmarked in this study. Results indicate that SoftHybrid and MSImpute achieve overall comparable performance; however, SoftHybrid demonstrates improved accuracy in abundance recovery, suggesting its advantage in quantitative analyses. Importantly, MSImpute relies on prior knowledge of sample groupings, which may not be readily available in many single-cell proteomics applications. Moreover, MSImpute was developed and validated primarily on bulk proteomics data with relatively low levels of missingness, whereas single-cell proteomics datasets are characterised by substantially higher sparsity and distinct data structures, potentially limiting its direct applicability in single-cell proteomics context.

Another modelling-based strategy is adopted in the QuantQC framework^25^, where missingness is explicitly modelled by integrating inferred biological annotations (e.g. cell types) and auxiliary variables such as cell size. While this design enables more interpretable modelling when such information is reliable, it introduces dependencies on upstream imputation, clustering, and annotation steps. In most single-cell proteomics datasets, where clustering may be unstable due to more biological noise and cell states often form continua rather than discrete groups, uncertainties introduced at early stages may propagate through the modelling process and reinforce patterns derived from imputed data. Additionally, QuantQC relies on external prior information for modelling missingness, such as cell size, which is not always available in single-cell sorting workflows (e.g., FACS-based sorting).

Rather than explicitly modelling missingness mechanisms, SoftHybrid uses data-driven proxies to approximate different missingness regimes within a continuous framework independent on external obtained information, and achieves a balanced representation, preserving the contrast between expressing and non-expressing cell types while avoiding systematic underestimation within expressing groups. This design provides a flexible and assumption-light alternative, particularly in settings where cell identities are ambiguous or difficult to define, and external information is limited. We see existing missingness modelling approaches as complementary to SoftHybrid, with explicit modelling frameworks offering greater interpretability when reliable annotations and cell sizes can be derived, and data-driven approaches such as SoftHybrid providing broader applicability in prior knowledge-limited settings.

Machine learning-based strategies have also gained increasing attention in proteomics imputation. However, these approaches require large sample sizes to perform reliably^50^. While this is feasible in transcriptomics, the lower throughput of proteomics makes it more challenging to generate datasets of comparable scale. Furthermore, these ML-based approaches have not yet been extensively validated on single-cell proteomics, where data sparsity may limit the effective training of complex models. In this context, SoftHybrid demonstrates consistent performance across both bulk and single-cell datasets and can be readily integrated into proteomics pipelines as a user-friendly R package. Furthermore, its improved performance at lower sample inputs and under DDA acquisition suggests that it may be applicable beyond single-cell proteomics. In particular, it may be beneficial for other sparse proteomics datasets characterised by high missingness, such as post-translational modification analyses, clinical samples, immune-enrichment workflows, or experiments performed on legacy mass spectrometers, where protein identifications are limited and missing values are more prevalent. While SoftHybrid shows favourable performance compared with existing approaches, several limitations should be considered. First, although SoftHybrid is not a machine learning-based method, it incorporates RF imputation as the MAR-oriented component. While RF provides strong predictive performance, it is computationally intensive, particularly for large datasets. Consequently, SoftHybrid may require considerable computational resources and longer runtimes in large-scale studies. For example, processing *Dataset SCP* (76 samples and 2,546 proteins) on a single CPU core required nearly 20 minutes to complete the full imputation workflow. Additionally, the weighting scheme is defined at the protein level and remain consistent across samples; therefore, it does not explicitly model sample-specific effects, including failed injections or variability in sample preparation (e.g. cell culture serum removal). RF may partially address this by leveraging global expression patterns, but such effects are not directly accounted for in the weighting framework. Finally, imputation is not universally appropriate and should be applied with caution^21^. As demonstrated in *Dataset HYE* (Figure 4G-H), although SoftHybrid reduces the magnitude of bias relative to other methods, deviations from the raw data, including both inflation and underestimation, can still occur. Similar patterns were observed in *Dataset Condition*, where all imputation methods introduced shifts in fold-change estimates compared to the raw data (Figure7D). These observations highlight that imputation may obscure meaningful biological signals, particularly in cases where missingness reflects true biological absence or condition-specific expression. Therefore, imputation should be applied selectively, for example in analyses that require complete data matrices. In other contexts, retaining the raw data may be preferable. Therefore, as with any imputation strategy, SoftHybrid does not substitute the need for careful experimental design and data acquisition. Improvements in instrumentation and acquisition methods^51^, which reduce missing values at the source remain essential for robust proteomics data analysis.

## Conclusion

Our results demonstrate that SoftHybrid provides a practical and broadly applicable approach for handling missing values in proteomics data. By balancing MAR- and MNAR-oriented imputations, it mitigates systematic biases and improves the recovery of biologically relevant patterns, supporting more robust downstream interpretation in single-cell settings. SoftHybrid is implemented as an R package and is available on GitHub, enabling straightforward application through a simple command-line interface.

## Associated content

### Data and code availability statement

The public *Dataset CellCycle* is available in ProteomeXchange Consortium via the PRIDE^52^ partner repository with the dataset identifier PXD024043, and the cell cycle marker list is available at https://github.com/theislab/singlecell_proteomics. The raw and processed mass spectrometry data of all three other in-house datasets have been deposited to the ProteomeXchange Consortium via the PRIDE^52^ partner repository with the dataset identifier PXD071819. The source data used for raw data interpretation and the R code used for analysis and benchmarking are available at https://github.com/YixinShiProteomics/SoftHybrid-paper. The open-source SoftHybrid package and its tutorials are freely available on GitHub at https://github.com/YixinShiProteomics/softHybridImpute.

### Supporting information

The following supporting information is available free of charge at ACS website http://pubs.acs.org

Supplementary Method design. Proof of missing rate and protein abundance as proxies for MNAR proportion

Figure S1. Overview of the end-to-end data pre-processing and analysis workflow for proteomics datasets.

Figure S2. Missing value distribution across different sample loads and acquisition modes in *Dataset HYE*.

Figure S3. Imputation methods are not neutral procedures

Figure S4. Influence of individual hyperparameters on the final weighting scheme.

Figure S5. Loess-based detection of elbow points across input amounts and acquisition modes in *Dataset HYE*.

Figure S6. SoftHybrid weighting distribution balancing MAR- and MNAR-oriented imputations across sample loads and acquisition modes in *Dataset HYE*.

Figure S7. Comparison of species-level ratio accuracy across imputation methods in DIA mode for *Dataset HYE*.

Figure S8. Comparison of species-level ratio accuracy across imputation methods in DDA mode for *Dataset HYE*.

Figure S9. Protein numbers involved in the benchmarking process across DIA and DDA datasets after raw and imputed processing at different sample amounts.

Figure S10. ROC curve comparison of imputation methods across input amounts and acquisition modes in *Dataset HYE*.

Figure S11. Comprehensive benchmarking of imputation methods across clustering accuracy, reproducibility, and biological interpretability.

Table S1. DeLong test of ROC analysis in comparisons of SoftHybrid vs other methods in DIA Table S2. DeLong test of ROC analysis in comparisons of SoftHybrid vs other methods in DDA

## Author Information

### Corresponding Author

**Roman Fischer** - Target Discovery Institute, Centre for Medicines Discovery, Nuffield Department of Medicine, University of Oxford, Roosevelt Drive, Oxford, OX3 7FZ, United Kingdom;

### Author contributions

Y.S conceived the idea, developed the algorithm, performed the single-cell sorting, mass spectrometry analysis, and bioinformatic analysis. S.D contributed to the data analysis and LC-MS/MS operation. S.D, P.D.C, S.T and R.F contributed to the experimental design. E.D and D.E contributed to cell culture and cell differentiation. G.B contributed to three-species sample preparation. R.F supervised this project. Y.S wrote the manuscript. All other authors provided feedback and revised the manuscript. All authors have given approval to the final version of the manuscript.

### Funding Sources

This work was supported by the Chinese Academy of Medical Sciences (CAMS) Innovation Fund for Medical Science (CIFMS), China (grant number: 2024-I2M-2-001-1).

### Notes

The authors declare no competing financial interest.

## For TOC Only

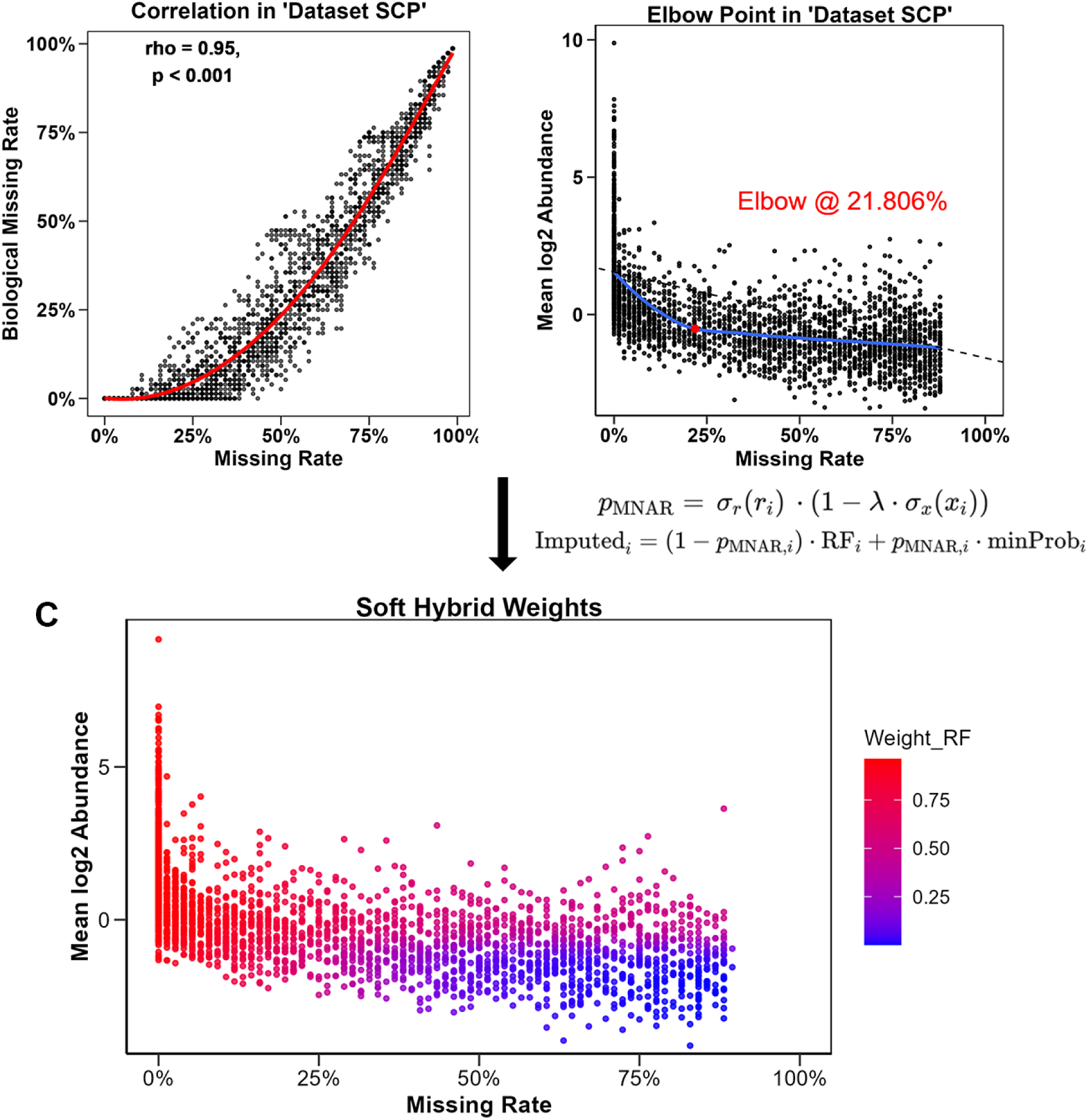

## Notes

### Competing Interest Statement

The authors have declared no competing interest.

### Summary of Updates

Updated algorithm optimisation, revised comparison with existing methods, and updated Figure 7.

