## Supplementary Figures for "SoftHybrid: A Hybrid Imputation Algorithm Optimised for Single-Cell Proteomics Data"

**Title of the primary article: SoftHybrid: A Hybrid Imputation Algorithm Optimised for Single-Cell Proteomics Data**

**Content tables:**

Supplementary Method design. Proof of missing rate and protein abundance as proxies for MNAR proportion

Figure S1. Overview of the end-to-end data pre-processing and analysis workflow for proteomics datasets.

Figure S2. Missing value distribution across different sample loads and acquisition modes in the Dataset HYE.

Figure S3. Imputation methods are not neutral procedures

Figure S4. Influence of individual hyperparameters on the final weighting scheme.

Figure S5. Loess-based detection of elbow points across input amounts and acquisition modes in the Dataset HYE.

Figure S6. SoftHybrid weighting distribution balancing MAR- and MNAR-oriented imputations across sample loads and acquisition modes in the Dataset HYE.

Figure S7. Comparison of species-level ratio accuracy across imputation methods in DIA mode for the Dataset HYE.

Figure S8. Comparison of species-level ratio accuracy across imputation methods in DDA mode for the Dataset HYE.

Figure S9. Protein numbers involved in the benchmarking process across DIA and DDA datasets after raw and imputed processing at different sample amounts.

Figure S10. ROC curve comparison of imputation methods across input amounts and acquisition modes in the Dataset HYE.

Figure S11. Comprehensive benchmarking of imputation methods across clustering accuracy, reproducibility, and biological interpretability.

Table S1. DeLong test of ROC analysis in comparisons of SoftHybrid vs other methods in DIA

Table S2. DeLong test of ROC analysis in comparisons of SoftHybrid vs other methods in DDA

***Supplementary Method*** ***Design***

***Missing rate and protein abundance as proxies for MNAR proportion***

In Dataset SCP of single-cell proteomics, the per-protein missing rate showed a significant negative correlation with mean ${log}_{2}$ protein abundance ($\rho$ = -0.545, *p*-value < 0.001), and a LOESS regression fit revealed a clear elbow point in this relationship; using a maximum-distance criterion, we identified the elbow at a 21.806% missing rate, suggesting a compositional shift in missingness around this threshold (Figure 3C).

To formalise the link between the missing rate and the missing-mechanism mixture, let $x_{i,j}$ be the abundance of protein $i$ in sample $j$. We model the MNAR probability as a monotone decreasing function $f(x_{i,j})$:

$\Pr\left( x_{i,j}= \phi| x_{i, j} \right)= f(x_{i,j})$	Eq.S1

For MAR, we assume a small, protein-specific constant probability $\rho_{i}^{MAR}$, typically $\ll1$

$\Pr\left( x_{i, j}= \phi\right)= \rho_{i}^{MAR}$	Eq.S2

Let $\lambda_{i} \in[0, 1]$ be the proportion of MNAR missingness for protein $i$. The overall per-entry missing probability is

$\Pr\left( x_{i, j}= \phi\right)= \lambda_{i}\cdot f\left( x_{i,j} \right)+(1-\lambda_{i})\cdot\rho_{i}^{MAR}$	Eq.S3

Averaging over samples yields the per-protein missing rate

$p_{i}= \mathbb{E}_{j}\left[ \Pr\left( x_{i, j}= \phi\right) \right]= \lambda_{i}\cdot\mathbb{E}_{j}\left[ f\left( x_{i,j} \right) \right]+(1-\lambda_{i})\cdot\rho_{i}^{MAR}$	Eq.S4

Differentiating with respect to $\lambda_{i}$ gives

$\frac{\partial p_{i}}{\partial\lambda_{i}}= \mathbb{E}_{j}\left[ f\left( x_{i,j} \right) \right]- \rho_{i}^{MAR}$	Eq.S5

In proteomics, $f(x)$ is often modelled as a sigmoid function around the MS detection threshold: $f\left( x \right) \to1$ for very low intensities and $f\left( x \right) \to0$ for high intensities. If protein intensities are approximately normal and centred near the threshold, then $\mathbb{E}\left[ f\left( x \right) \right]\approx0.5$, which in practice exceeds $\rho_{i}^{MAR}$. Therefore,

$\mathbb{E}\left[ f(x) \right] > \rho_{i}^{MAR} \Rightarrow\frac{\partial p_{i}}{\partial\lambda_{i}} >0$	Eq.S6

Eq.S6 demonstrates that more MNAR dominance ($\lambda_{i}$ increases) increases the missing rate ($p_{i}$ increases).

To validate this, we applied **BIND** developed by Guo et al., to classify biological (MNAR-like) and technical (MAR-like) missingness. The predicted biological missing rate correlated strongly with the overall missing rate ($\rho$ = 0.95, *p*-value < 0.001) and showed a sharp transition near the 25% threshold, consistent with the LOESS elbow (Figure 3A).

***Supplementary Figures***

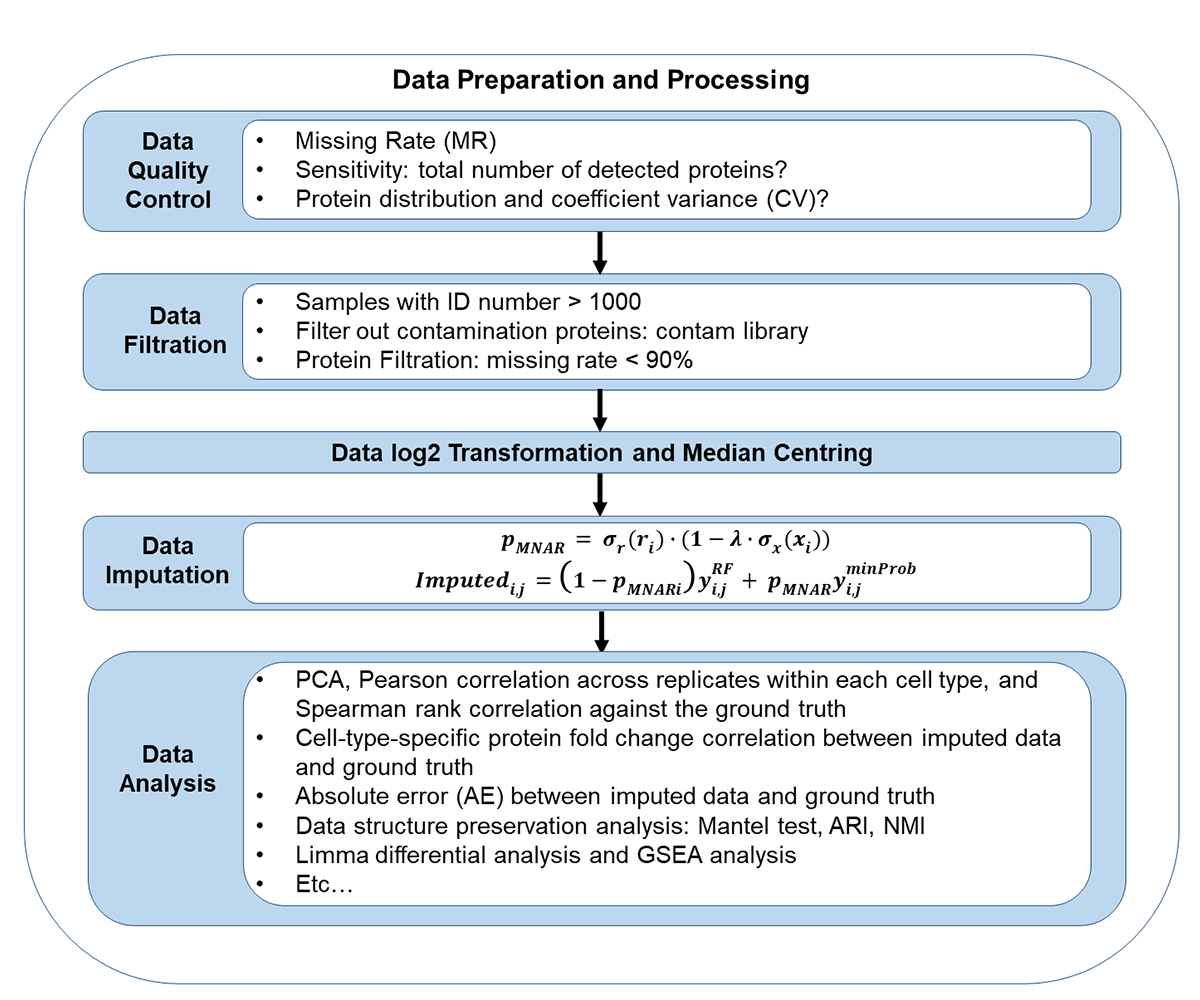

**Figure S1. Overview of the end-to-end data pre-processing and analysis workflow for proteomics datasets.** The pipeline consists of sequential steps including data quality control, filtration, transformation, and median-centred normalisation, followed by SoftHybrid-based imputation and downstream analyses. Quality control assesses the missing rate, sensitivity, and consistency of protein distribution. Filtration removes low-quality and contaminant proteins, while imputation adaptively balances MNAR and MAR assumptions through the SoftHybrid weighting formula. Downstream analyses include PCA, replicate correlations, absolute error calculation, data structure preservation analysis, differential expression analysis (limma), and GSEA-based biological interpretation.

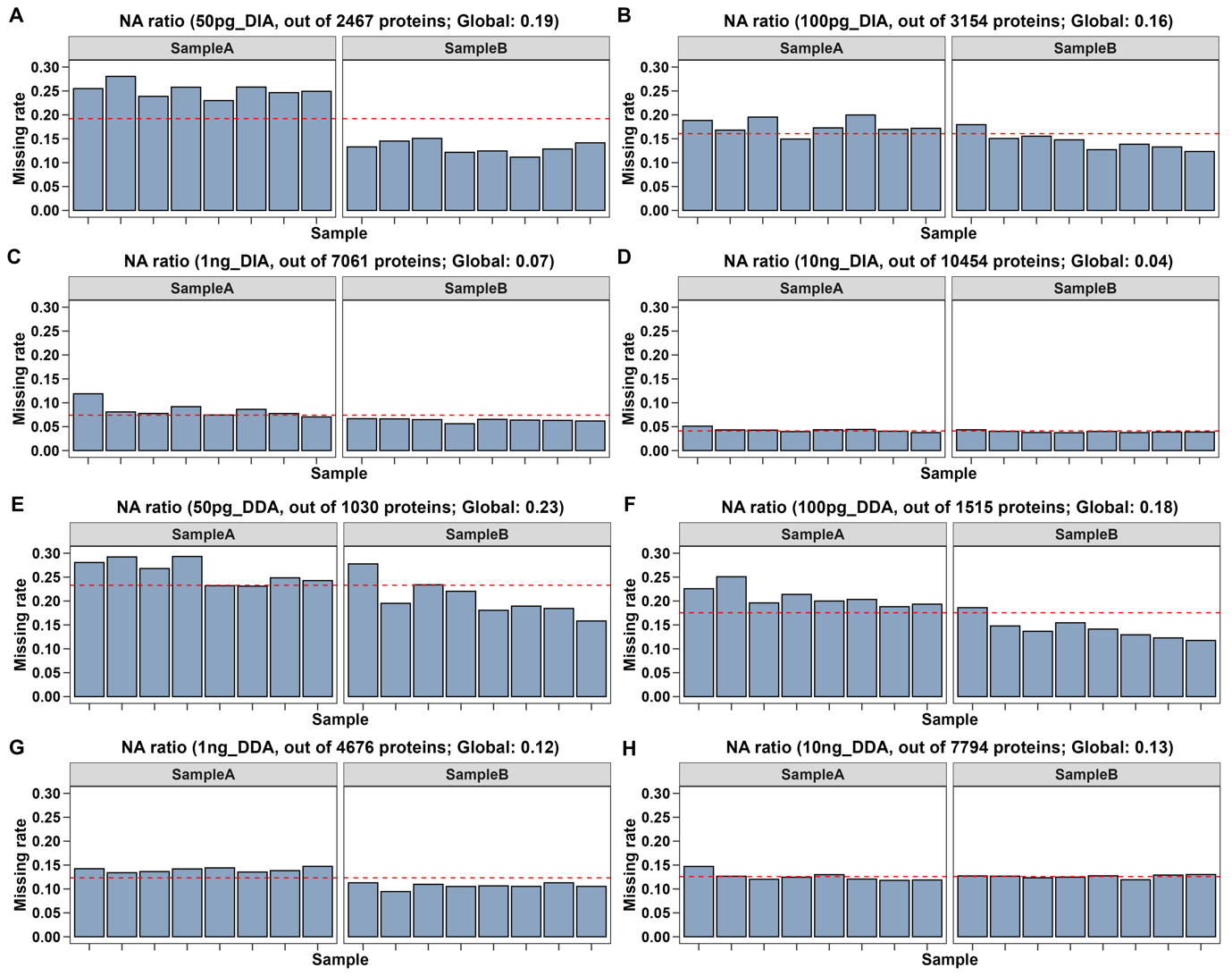

**Figure S2. Missing value distribution across different sample loads and acquisition modes in the Dataset HYE.** Bar plots show per-sample missing rates under different input amounts and acquisition modes (50 pg to 10 ng; DIA: A-D; DDA: E-H). Sample A and Sample B correspond to mixtures with distinct species proportions. The global missing rate for each condition is indicated by a red dashed line. Lower sample amounts exhibit higher missing rates, particularly under DDA acquisition, reflecting increased stochasticity and reduced identification sensitivity at low input levels.

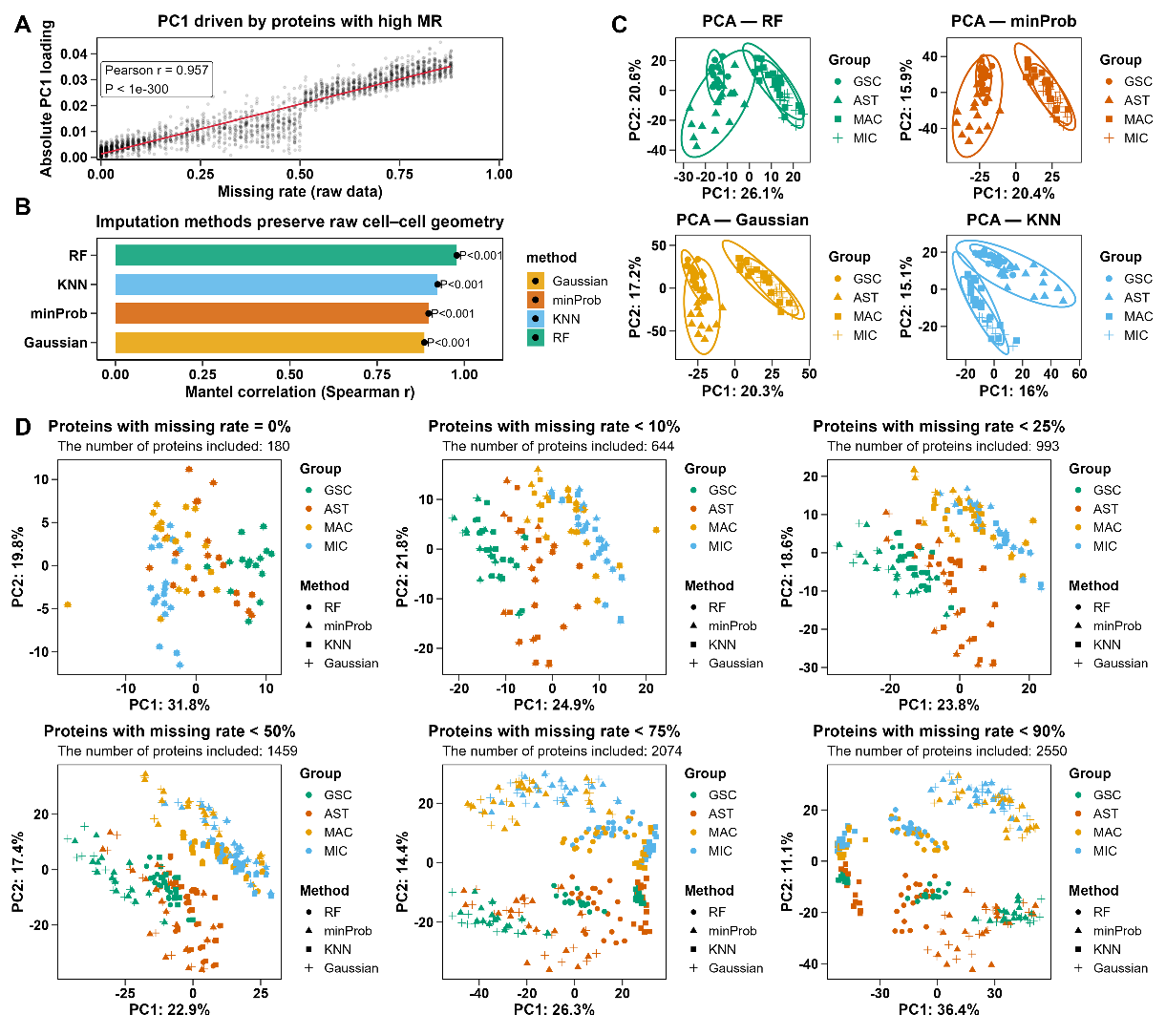

**Figure S3. Imputation methods are not neutral procedures.** (A) Relationship between protein missing rate in the raw single-cell proteomics dataset and the absolute loading of the first principal component (PC1). Each point represents a protein. Proteins with higher missing rates contribute more strongly to PC1, indicating that the primary variance captured by PCA is strongly driven by missingness patterns in the raw data (Pearson *r* = 0.957, *p* < 1 × 10-300). (**B**) Preservation of raw cell-cell geometry after imputation. For each imputation method, the cell-cell distance matrix derived from the imputed datasets was compared with the raw dataset using a Mantel test. All imputation methods significantly preserved the underlying cellular distance structure (Spearman Mantel correlation, 999 permutations, r > 0.75, *p* = 0.001. (**C**) Separate PCA plots for different imputation methods on the same raw data matrix. (**D**) PCA separation becomes increasingly driven by imputation methods as protein missingness increases.

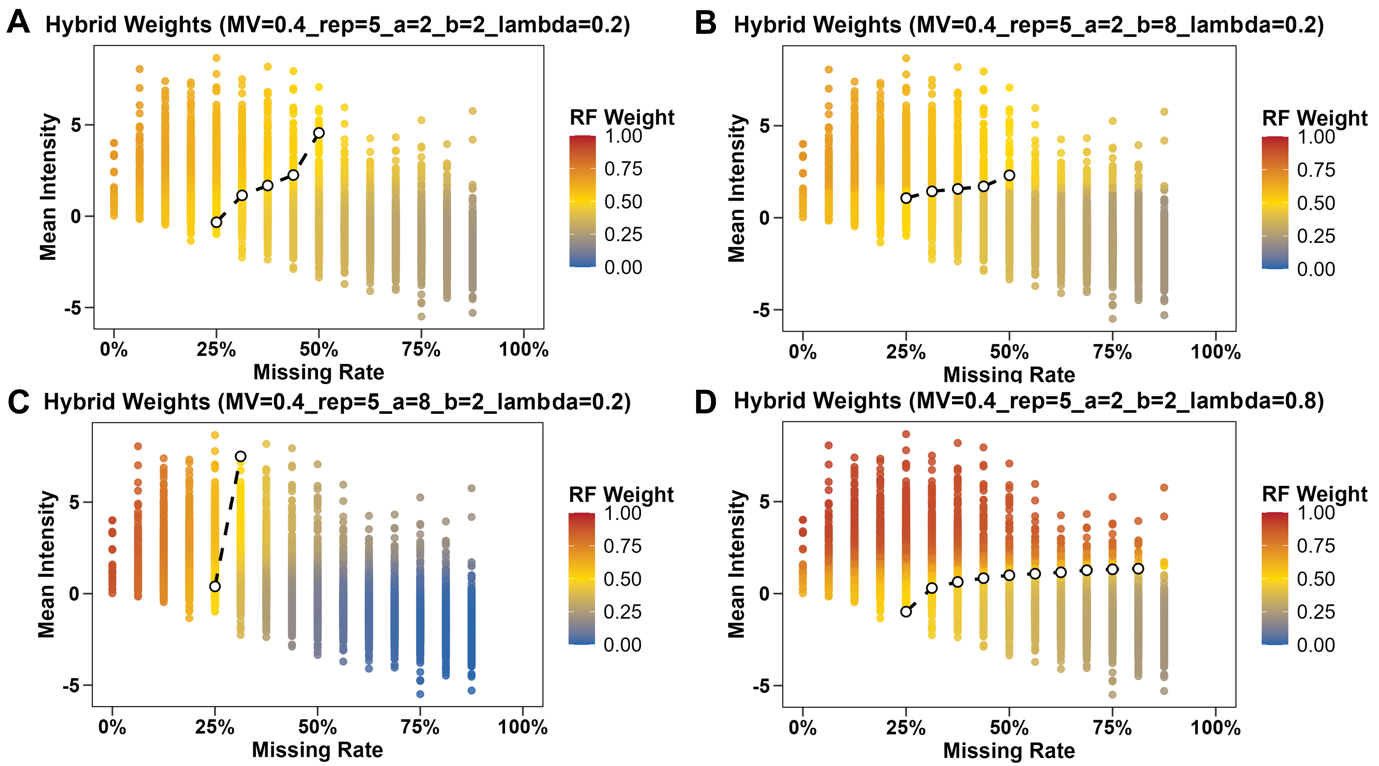

**Figure S4. Influence of individual hyperparameters on the final weighting scheme.** RF weights in the hybrid imputation model are shown as a colour gradient from blue (0) to red (1) under different combinations of hyperparameters. Each point represents a protein, position by its missing rate (x-axis) and mean intensity (y-axis). Panels A-D illustrate the effects of varying parameters a, b, and $\lambda$ on the resulting RF weight distribution. Here, a controls the steepness of the missing-rate-dependent transition ($a_{r}$) and b controls the intensity-dependent component ($a_{x}$). White circle indicate the transition boundary where RF weight = 0.5, separating regions dominated by RF-based and minProb-based imputation.

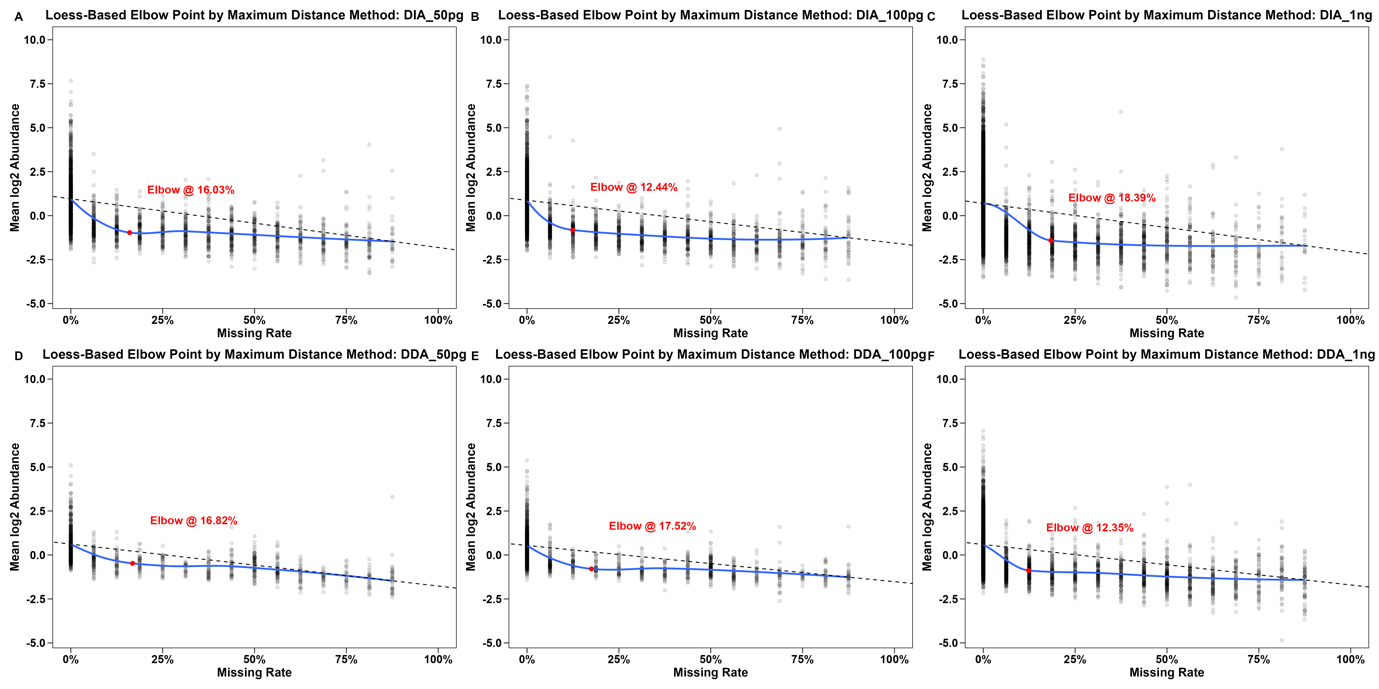

**Figure S5. Loess-based detection of elbow points across input amounts and acquisition modes in the Dataset HYE.** Scatter plots show the relationship between mean log₂ protein abundance and missing rate under different sample loads (50 pg to 1 ng) and acquisition modes (DIA: A-C; DDA: D-F). The blue line represents the loess-fitted curve, and the red dot indicates the elbow point determined by the maximum distance method. The detected elbow points (12-19%) mark the transition between proteins with biologically meaningful signal and those dominated by missingness noise, providing adaptive thresholds for SoftHybrid weighting.

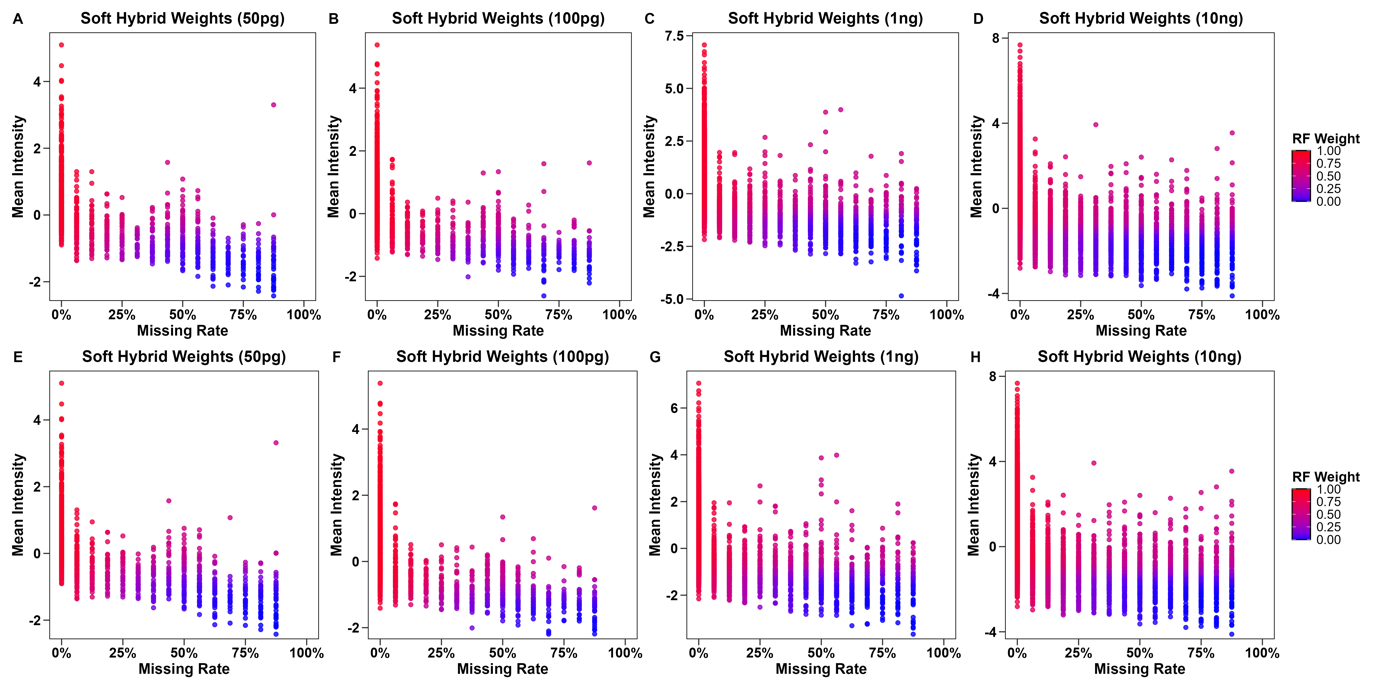

**Figure S6. SoftHybrid weighting distribution balancing MAR- and MNAR-oriented imputations across sample loads and acquisition modes in the Dataset HYE.** Scatter plots display the adaptive weighting (RF weight, colour-coded) applied by the SoftHybrid algorithm for each protein according to its mean intensity and missing rate across varying input amounts (50 pg to 10 ng) and acquisition modes (DIA: A-D; DDA: E-H). Proteins with higher abundance and lower missing rate receive greater Random Forest (MAR) weights, while low-abundance, high-missing-rate proteins are predominantly imputed by the minProb (MNAR) component, reflecting SoftHybrid’s continuous balancing between MAR and MNAR assumptions.

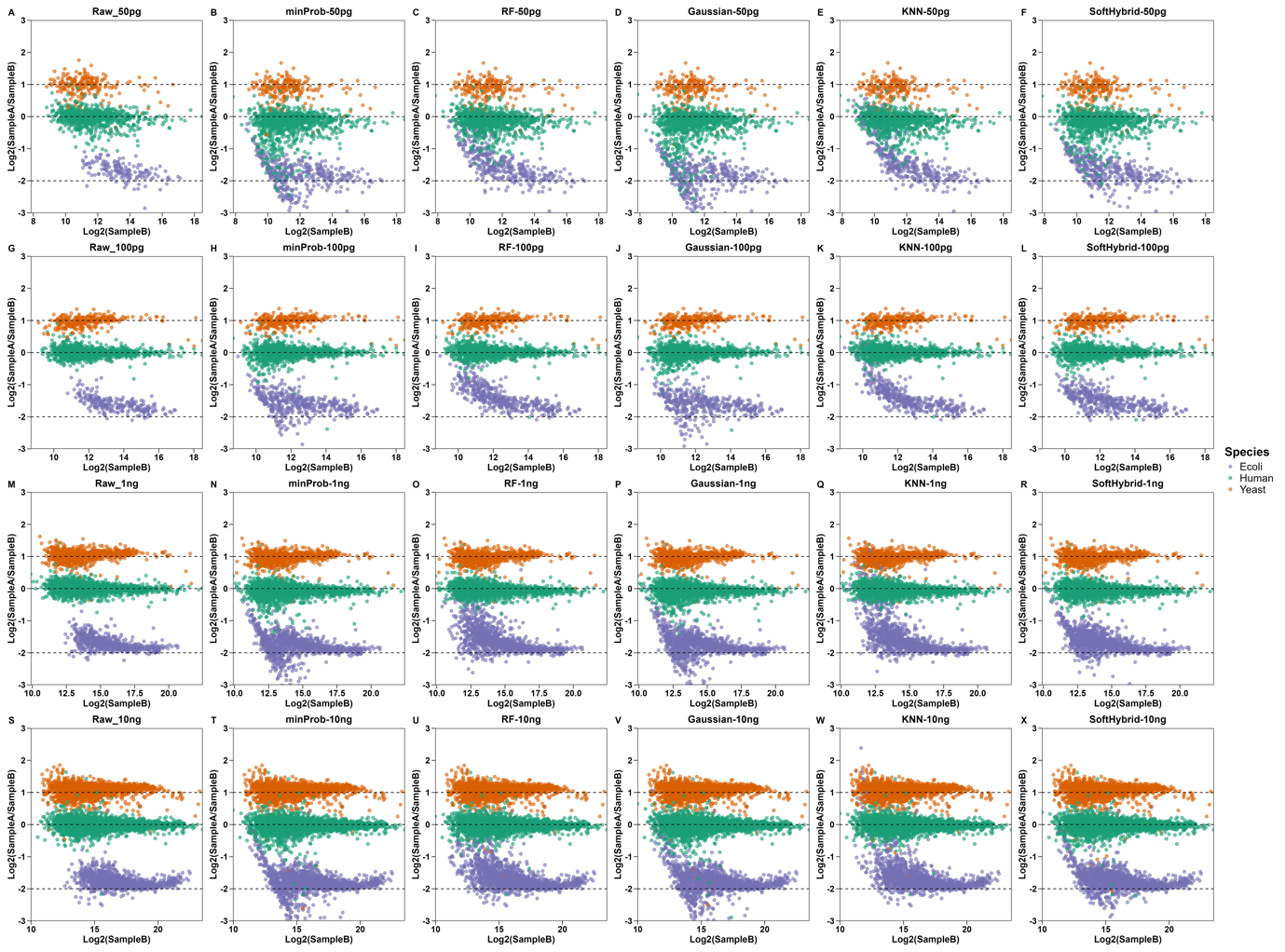

**Figure S7. Comparison of species-level ratio accuracy across imputation methods in DIA mode for the Dataset HYE.** Scatter plots show log₂(Sample A/Sample B) versus log₂(Sample B) across different input amounts (50 pg to 10 ng) and imputation strategies (Raw, minProb, RF, Gaussian, KNN, and SoftHybrid). Each point represents a quantified protein, coloured by species (E. coli, Yeast, Human). While traditional MNAR-based methods (e.g., minProb) introduce systematic underestimation in mid-intensity ranges and MAR-based methods (e.g., RF) overestimate protein abundances at the low end, SoftHybrid minimises bias across abundance scales, preserving expected species ratios and maintaining quantitative accuracy even at low input levels.

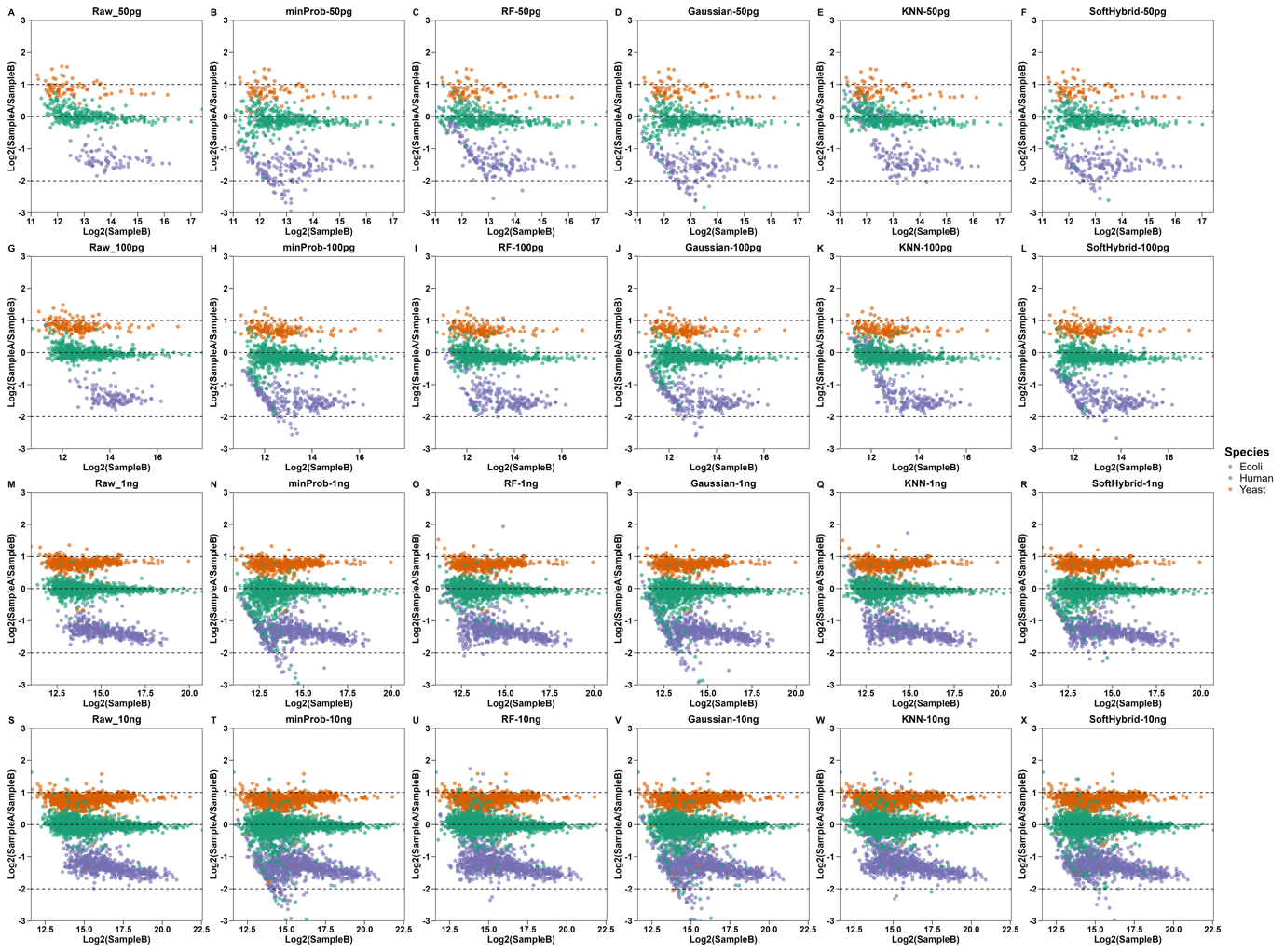

**Figure S8. Comparison of species-level ratio accuracy across imputation methods in DDA mode for the Dataset HYE.** Scatter plots show log₂(Sample A/Sample B) versus log₂(Sample B) across different input amounts (50 pg to 10 ng) and imputation strategies (Raw, minProb, RF, Gaussian, KNN, and SoftHybrid). Each point represents a quantified protein, coloured by species (E. coli, Yeast, Human). Compared with DIA acquisitions, DDA mode exhibits higher stochastic missingness and wider variance. SoftHybrid effectively reduces systematic misestimation and improves ratio accuracy across species, demonstrating enhanced robustness at lower sampling depths in DDA data.

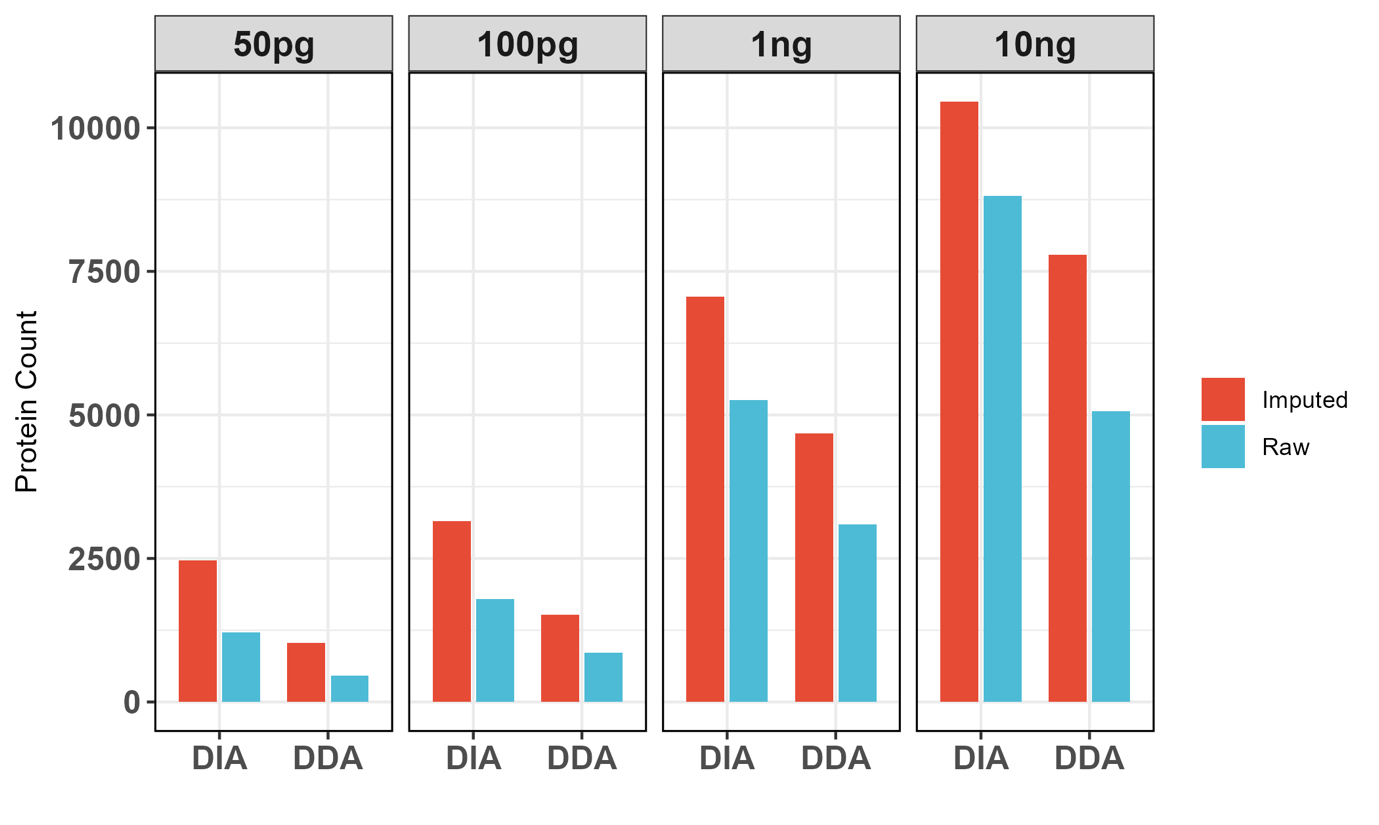

**Figure S9. Protein numbers involved in the benchmarking process across DIA and DDA datasets after raw and imputed processing at different sample amounts.** Bar plots show the number of identified proteins for DIA and DDA at 50 pg, 100 pg, 1 ng, and 10 ng, with raw data (blue) and imputed data (red) displayed side-by-side. Within each dose, DIA is shown on the left and DDA on the right. Imputation consistently increases the number of quantified proteins across all acquisition modes and sample amounts, with the largest gains observed at the lowest input levels.

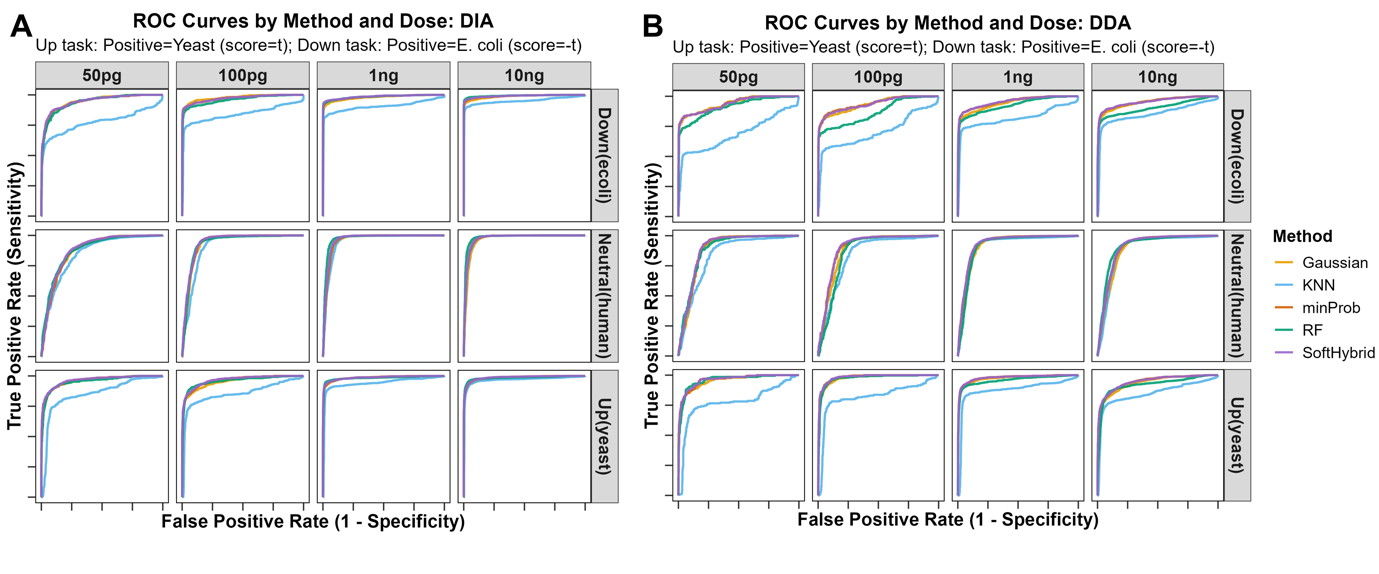

**Figure S10. ROC curve comparison of imputation methods across input amounts and acquisition modes in the Dataset HYE.** ROC curves illustrate the classification performance for upregulated (Yeast), neutral (Human), and downregulated (E. coli) species across different sample loads (50 pg-10 ng) and acquisition modes (A, DIA; B, DDA). Each curve represents a distinct imputation method (Gaussian, KNN, minProb, RF, SoftHybrid). SoftHybrid consistently achieves higher sensitivity and specificity, particularly under low-input conditions, indicating improved recovery of true biological differences while maintaining low false-positive rates.

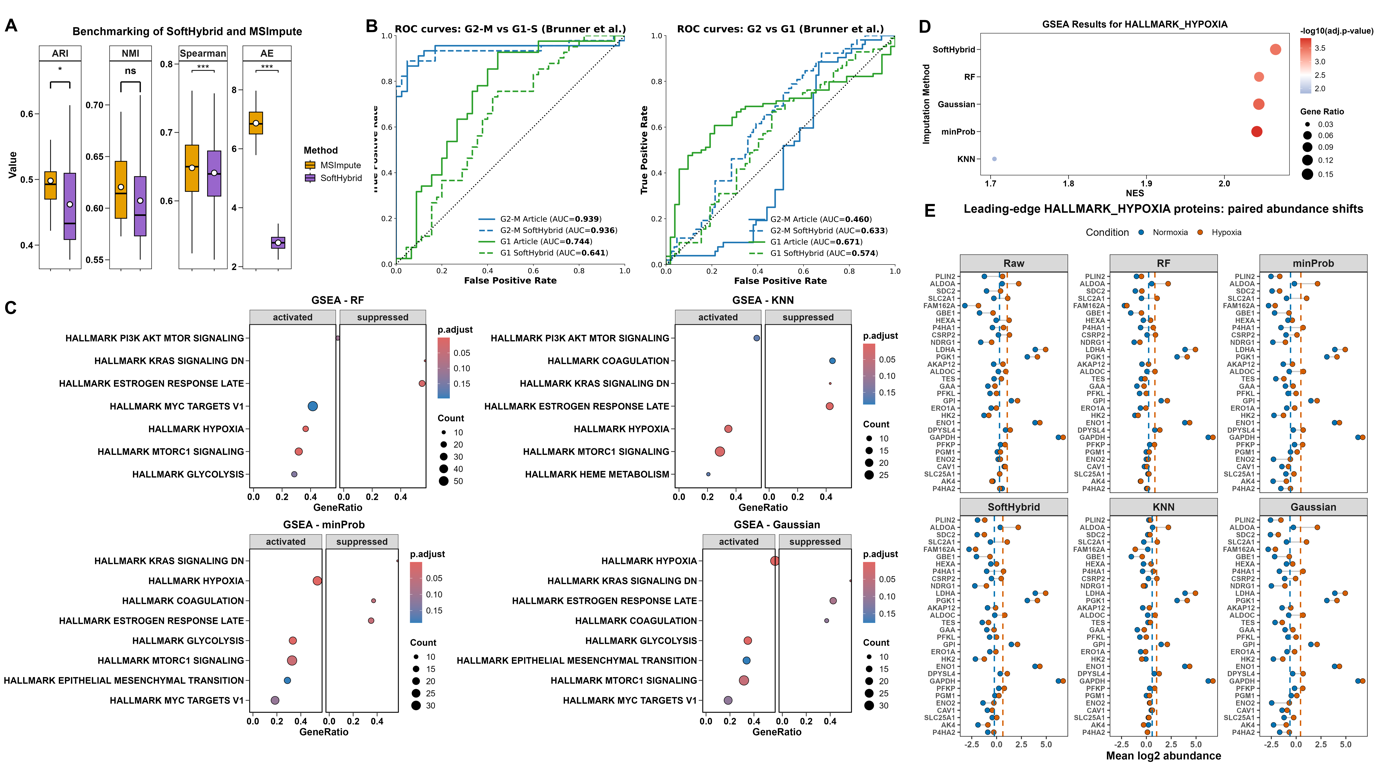

**Figure S11. Comprehensive benchmarking of imputation methods across clustering accuracy, reproducibility, and biological interpretability.** (A) ARI, NMI, Spearman correlation and absolute error (AE) comparison between SoftHybrid and MSImpute in Dataset SCP. (B) ROC AUC comparison of SoftHybrid and zero imputation in Dataset CellCycle using G2M marker-based and G1 marker-based scoring to distinguish between G2-M vs G1-S stages (AUC = 0.936 vs 0.939, *p* = 0.885 for G2-M marker scoring; AUC = 0.641 vs 0.744, *p* = 0.079 for G1 marker scoring) and G2 vs G1 stages (AUC = 0.633 vs 0.540, *p* = 0.321 for G2-M marker scoring; AUC = 0.574 vs 0.671, *p* = 0.321 for G1 marker scoring). (C) GSEA results of differential analysis (hypoxia vs. normoxia) for representative methods in the Dataset Condition. (D) GSEA results for HALLMARK HYPOXIA across different imputation methods (SoftHybrid, RF, Gaussian, KNN, minProb). (E) Paired protein abundance shifts (fold changes) for individual leading edge HALLMARK HYPOXIA proteins between normoxia and hypoxia conditions, across different imputation methods (SoftHybrid, RF, Gaussian, KNN, minProb) and raw data.
